# Overventilation-induced airspace acidification increases susceptibility to Pseudomonas pneumonia

**DOI:** 10.1101/2024.08.07.603041

**Authors:** Matthias Felten, Chunjiang Tan, Sebastian Ferencik, Jingjing Li, Eleftheria Letsiou, Jasmin Lienau, Holger Müller-Redetzky, Andreas Hocke, Theresa C. Brömel, Achim D. Gruber, Gopinath Krishnamoorthy, Matthias Ochs, Christina Brandenberger, Qi Zhang, Wolfgang M. Kuebler, Martin Witzenrath

## Abstract

Ventilator-associated pneumonia (VAP) is the most frequent nosocomial infection in critically ill patients. Local pH variations affect bacterial growth. Whether airway acidification contributes to the pathogenesis and pathophysiology of *Pseudomonas aeruginosa* (PA)-induced VAP is currently unknown. This study was undertaken to investigate the role and mechanisms of airspace acidification by mechanical ventilation (MV) in PA-induced VAP.

C57BL/6J mice were subjected to high (HVt: 34 mL/kg) or low (LVt: 9 mL/kg) tidal volume MV for 4 h. PA was instilled *via* the tracheal tube, and animals were allowed to recover from sedation and breathe spontaneously for 24 h following extubation. Fluorescence microscopy was applied to determine alveolar pH in ex vivo perfused and ventilated murine lungs. Bacterial growth and adhesion on cyclically stretched A549 and human alveolar epithelial cells was examined.

Upon PA infection, HVt mice showed increased alveolo-capillary permeability, elevated lung and blood leukocyte counts, and higher bacterial load in lungs and extrapulmonary organs as compared to LVt controls. HVt MV induced acidification of alveolar lining fluid (ALF) in lungs and decreased pulmonary expression of Na^+^/H^+^ exchanger 1 (NHE1). Inhibition of NHE1 enhanced PA growth *in vitro* on alveolar epithelial cells and increased pulmonary bacterial loads in LVt-MV mice *in vivo*.

In a novel murine VAP model, key characteristics of PA-VAP were replicated. HVt MV induced mild VILI with acidification of airway lining fluid, increasing susceptibility to PA pneumonia. NHE1 was identified as critical factor for MV-induced airspace acidification, and thus as potential target to combat PA-VAP.

## Introduction

Ventilator-associated pneumonia (VAP) is among the most common intensive care unit-acquired infections (1) and associated with prolonged hospital stay, high treatment costs and increased mortality (2). Mechanical ventilation (MV) is a life-saving treatment but prolonged duration of MV increases the susceptibility of patients to pneumonia by approximately 6- to 20-fold when compared to non-ventilated patients (3). In injured lungs with non-ventilated areas due to edema, cellular infiltrates and/or atelectasis, even “protective” MV may result in biotrauma due to overdistension of the residual ventilated lung areas, a process termed ventilator-induced lung injury (VILI).

*Pseudomonas aeruginosa* (PA) is frequently identified as etiological agent of VAP (4). Recurrence and mortality rate in PA-associated VAP remain a critical challenge for clinical management that is aggravated by the lack of reliable prognostic clinical markers of VAP as well as the increasing emergence of multidrug-resistant PA isolates (5, 6). Understanding the physiological and immunological responses to MV and establishing their impact on bacterial infection and disease progression is critical for identifying measures to reduce the burden of VAP.

Host susceptibility and pathogen virulence essentially dictate the manifestation of infectious diseases. Interestingly, airway acidification is commonly detected in patients with acute lung injury (ALI) and acute respiratory distress syndrome (ARDS) (7, 8). Inflammatory processes may cause local acidification, resulting in physicochemical imbalance that directly augments the fitness of bacterial pathogens (9, 10), as previously reported in a porcine cystic fibrosis model (11). The cystic fibrosis transmembrane conductance receptor (CFTR) mediated bicarbonate secretion as well as the Na+ / H+ Exchanger 1 (NHE1) gated proton secretion may affect extracellular pH regulations (12) and both are known to be sensible to mechanical stress under different conditions (13, 14).

In consequence, we hypothesized that mechanical overventilation itself may alter the alveolar pH and thereby significantly impact host susceptibility to pursue pneumonia. Yet, in many animal models of VAP, bacterial infection is established prior to the onset of MV, contrasting the clinical sequence of events (15–17). As such, animal models of infection subsequent to MV may prove more appropriate to investigate the mechanistic links between VILI and VAP progression and severity. Such an approach has previously been successfully established in rat and porcine models (18, 19), but not yet in mice despite the obvious advantages regarding availability of molecular tools and genetically altered strains for mechanistic analyses (20).

In this study, we demonstrate that MV induces airspace acidification which in turn promotes PA adhesion and infection. Using our newly established murine PA-VAP model by mechanically ventilating mice prior to endotracheal instillation of PA, we show that mildly injurious overventilation renders mice more susceptible to PA pneumonia. Using this model in combination with ex vivo perfused and ventilated murine lungs and cyclically stretched human lung epithelial cells, we identified NHE1 as important mediator of MV-induced airway acidification. Some of the results of these studies have been previously reported in the form of an abstracts.

## Materials and methods

Animal experiments were approved by local and governmental authorities and carried out in accordance with EU-Directive 2010/63/EU.

### Murine VAP

Female C57BL/6J mice (8-10 weeks) were anaesthetized, orotracheally intubated and mechanically ventilated (FlexiVent^TM^, Montreal, Canada) for 4h with high (HVt: 34 mL/kg) or low tidal volume (LVt: 9 mL/kg) (21) or not ventilated (NV). For infection, 5×10^4^ colony-forming units (CFU) PA103 (ATCC #29260) or phosphate-buffered saline (PBS) was instilled *via* the endotracheal tube. After post-anesthesia recovery, mice were extubated, and breathed spontaneously for 24 h. Procedures of cell isolation, bacterial load enumeration, and histology are detailed in the online supplement. For treatment experiments, 3 mg/kg NHE1 inhibitor cariporide solved in DMSO were injected i.v. via the tail vein after narcosis.

### *Ex vivo* isolated perfused mouse lung

Lungs were harvested, perfused, and ventilated with HVt (20 mL/kg) or LVt (9 mL/kg) for 2h (22). pH-sensing fluorescent probe pHrodo™ Red Dextran (Thermo Fisher Scientific, Waltham, USA) was endotracheally instilled*. S*ubpleural alveoli were imaged *in situ* by confocal fluorescence microscopy.

### *In vitro* cell stretch and infection

Human A549 cells (ATCC #CCL-185) and human primary alveolar epithelial cells (hpAEC) were subjected to cyclic stretch (CS: elongation 18%, 20 cycles/min) or kept static (NCS) for 24h using the FlexCell^TM^. GFP-PAO1 (laboratory strain) attachment to cells was determined by immunofluorescence microscopy. PAO1 growth was assessed in acidified, untreated or alkalized RPMI.

### Data analysis

Statistical analysis was performed using the GraphPad Prism 9.30 (San Diego, CA, USA) software. P values < 0.05 were considered statistically significant.

## Results

### High tidal volume ventilation (HVt) increases the susceptibility to *P. aeruginosa* infection

To model the clinical sequence of PA-induced VAP, mice were subjected to MV with either high (HVt 34 mL/kg) or low tidal volume (LVt 9 mL/kg) followed by infection with PA, or PBS as uninfected control (**figure 1A**). Non-ventilated mice served as control. To assess the impact of HVt vs. LVt, changes in lung mechanics (**figure 1B**) were assessed over the initial 4 h of MV. Contrasting stable lung mechanics in LVt, mean airway pressure (MAP) gradually increased in HVt. Further, dynamic elastance, static compliance, and inspiratory capacity (IC) substantially increased exclusively due to HVt, demonstrating development of lung injury (**figure 1B; supplementary figure E1**).

**Figure 1:**
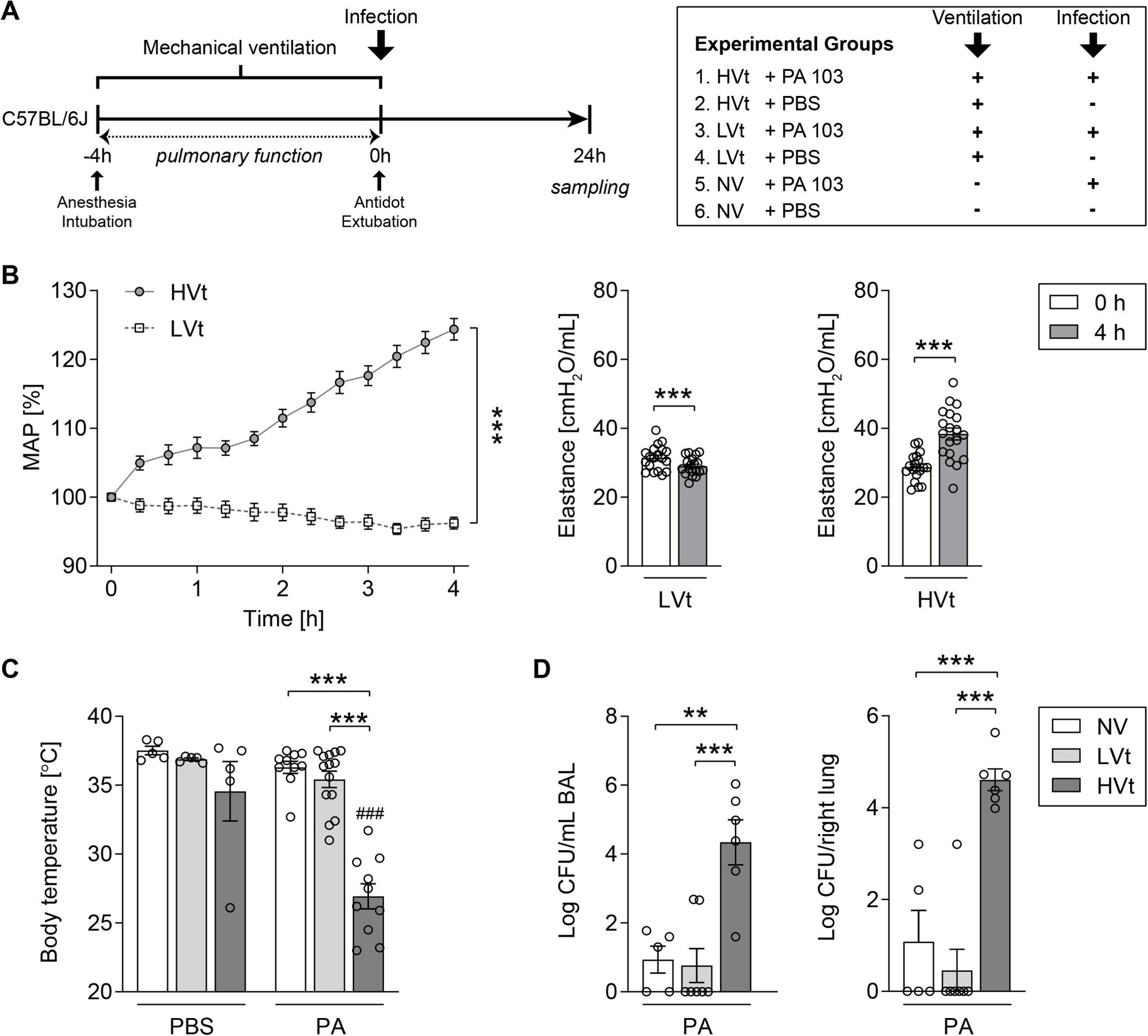
High tidal volume ventilation increases the susceptibility to *P. aeruginosa* infection. Mice were mechanically ventilated with HVt (34 mL/kg), LVt (9 mL/kg) for 4h or ventilated for 10 min (LVt) as NV control. After MV either PA103 or sterile PBS was instilled intratracheally and mice were extubated and left breathing spontaneously for 24h. (**A**) Experimental design of the *in vivo* experiments. (**B**) Mean airway pressure (MAP) was continuously measured during 4h MV, and dynamic elastance was measured *via* recruitment maneuvers at the beginning (0 h) and end of MV (4h). (**C**) Body temperature was measured at the end point of experiment. (**D**) Pulmonary bacterial colony-forming units (CFUs) in BAL and lung homogenates were determined. In (**A**), n = 19 per group; in (**C**), n = 5 (PBS) or n = 10-14 (PA); in (**D**), n = 5-7 per group. Data are presented as mean ± SEM, and analyzed using two-way repeated measures ANOVA (**A** left), Wilcoxon signed ranked test (**B** middle, right), two-way (**C**) or one-way (**D**) ANOVA and Sidak’s multiple comparisons test. ^#^ indicates significant difference compared with respective sham-infected group, * as indicated. \*\**p*<0.01, ***^/###^*p*<0.001. *BAL,* bronchoalveolar lavage; *HVt,* high tidal volume; *LVt,* low tidal volume; *MV,* mechanical ventilation; *NV*, non-ventilated; *PA, Pseudomonas aeruginosa*; *PBS,* phosphate-buffered saline.

Next, we examined pneumonia severity after MV. Body temperature and pulmonary bacterial loads were assessed 24 h post infection. Compared to other groups, body temperature declined in mice subjected to HVt and PA infection (**figure 1c**), and pulmonary bacterial growth in the HVt-PA group exceeded pulmonary bacterial growth in LVt or NV mice approximately 4-fold (**figure 1D**). These findings demonstrate significantly increased bacterial burden and disease severity upon PA infection after HVt ventilation, as compared to LVt and NV mice, suggesting higher susceptibility to PA infection in HVt mice.

Further, we investigated whether HVt ventilation impacts the inflammatory response to subsequent PA infection. Analysis of inflammatory cells in BALF and lungs revealed a trend towards increased PMN counts in all infected groups as compared to controls; yet, differences only reached significance in the HVt groups. (**figure 2A**). Multiplex immunoassays showed marked elevation of inflammatory cytokines and chemokines (IL-6, IL-10, CXCL-1, MCP-1, CXCL-5, IL-1α, MIP-1α, MIP-1β, eotaxin) in BALF of HVt-PA mice as compared to infected LVt and NV mice (**figure 2B, supplementary figure E2A**). IL-1ß concentrations were similarly increased between HVt compared to NV, but not versus LVt group. These results were corroborated by gene expression analysis in lung tissue (**supplementary figure E2B**). Likewise, alveolo-capillary permeability increased in mice subjected to HVt and PA infection as compared to other infected groups (**figure 2C**), suggesting that HVt MV led to an increased inflammatory response and consequently lung barrier disruption upon PA infection.

**Figure 2:**
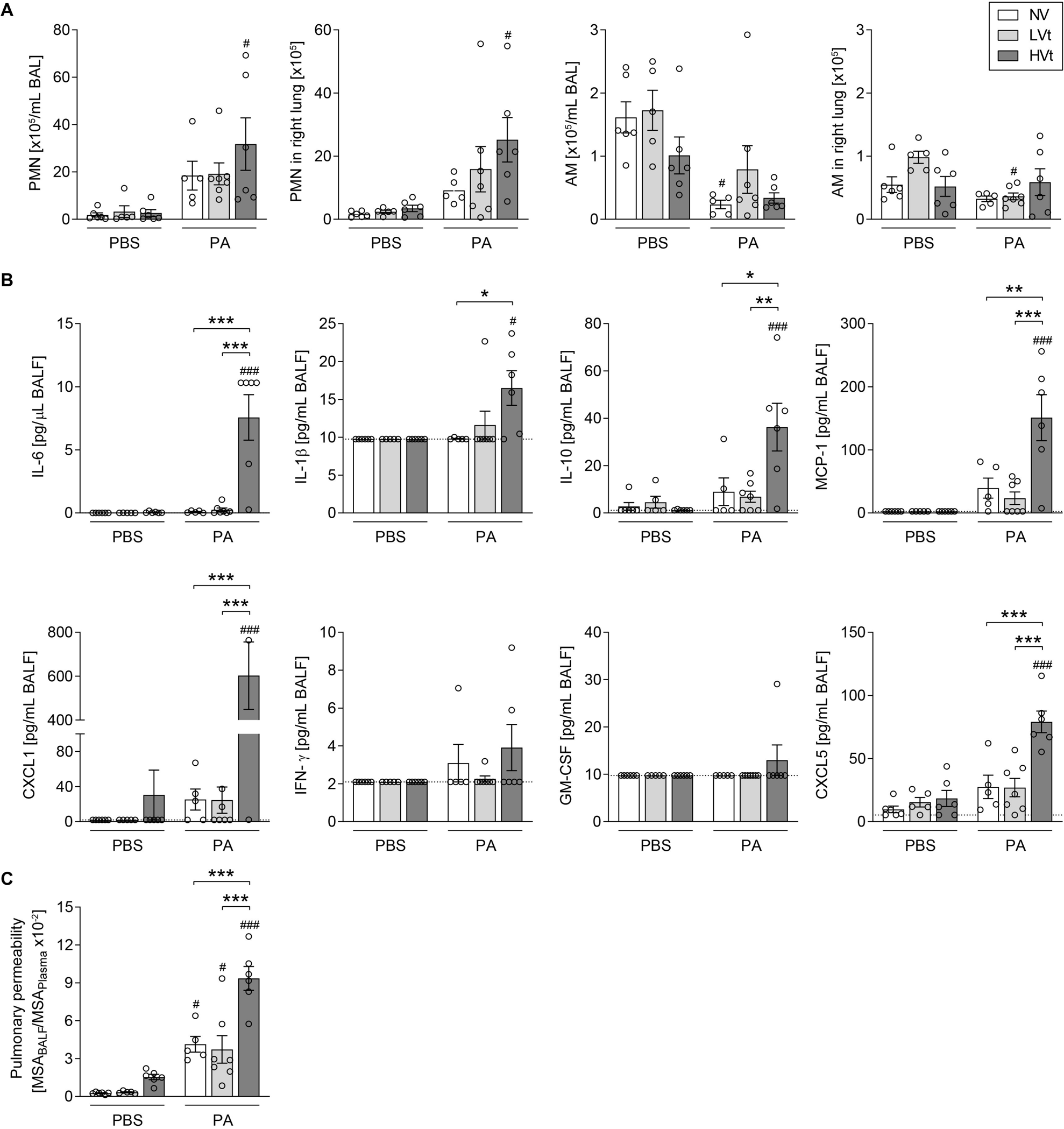
Pulmonary inflammation is increased in *P. aeruginosa* pneumonia subsequent to high tidal volume ventilation. Mice were mechanically ventilated with HVt (34 mL/kg), LVt (9 mL/kg) for 4h or ventilated for 10 min (LVt) as NV control. After MV either PA103 or sterile PBS was instilled intratracheally and mice were extubated and left breathing spontaneously for 24h. (**A**) Numbers of PMNs and alveolar macrophages in BAL and lung homogenates were differentiated and quantified *via* flow cytometry. (**B**) Inflammatory cytokines and chemokines in BALF were measured with multiplex beads-based immunoassay technique. (**C**) Alveolar-capillary permeability was assessed by measuring albumin in BALF and plasma and calculating the ratio. Data are presented as mean ± SEM; n = 5-6 (PBS) or n = 5-7 (PA). In (**B**), *dotted lines* indicate lower detection limits. ^#^ indicates significant difference compared with respective sham-infected group, * indicates significant difference between different ventilation regimes (as indicated). *^/#^*p*<0.05, \*\**p*<0.01, ***^/###^*p*<0.001 (two-way ANOVA and Sidak’s multiple comparisons test). *AM*, alveolar macrophages; *BAL,* bronchoalveolar lavage; *BALF,* bronchoalveolar lavage fluid; *CXCL*, C-X-C motif chemokine ligand; *GM-CSF*, granulocyte-macrophage colony-stimulating factor*; HVt,* high tidal volume; *IFN*, interferon; *IL*, interleukin; *LVt,* low tidal volume; *MCP*, monocyte chemoattractant protein; *MSA*, mouse serum albumin; *MV,* mechanical ventilation; *NV*, non-ventilated; *PA, Pseudomonas aeruginosa*; *PBS,* phosphate-buffered saline; *PMN*, polymorphonuclear cells.

### HVt promotes systemic dissemination of *P. aeruginosa*

Next, we examined the impact of MV on extra-pulmonary dissemination of PA. Remarkably, bacterial isolation from blood, liver and spleen revealed severe dissemination of PA infection in HVt mice, yet not in LVt or NV mice (**figure 3A**). Similar to pulmonary leukocyte profiles, blood PMN counts were significantly higher in HVt-PA mice as compared to other groups (**figure 3B**). Further, plasma levels of IL-6, CXCL-1, TNF-α, and MCP-1 were markedly increased only in HVt-PA infected mice suggesting the activation of acute inflammatory responses to bacterial dissemination (**figure 3C**). To probe for extrapulmonary organ damage due to systemic PA infection, we assessed markers of liver and kidney function. In HVt-PA mice, a trend to elevated AST levels (p=0.0535) and markedly increased total bilirubin, triglyceride, and urea concentration in plasma were detected (**supplementary figure E3**).

**Figure 3:**
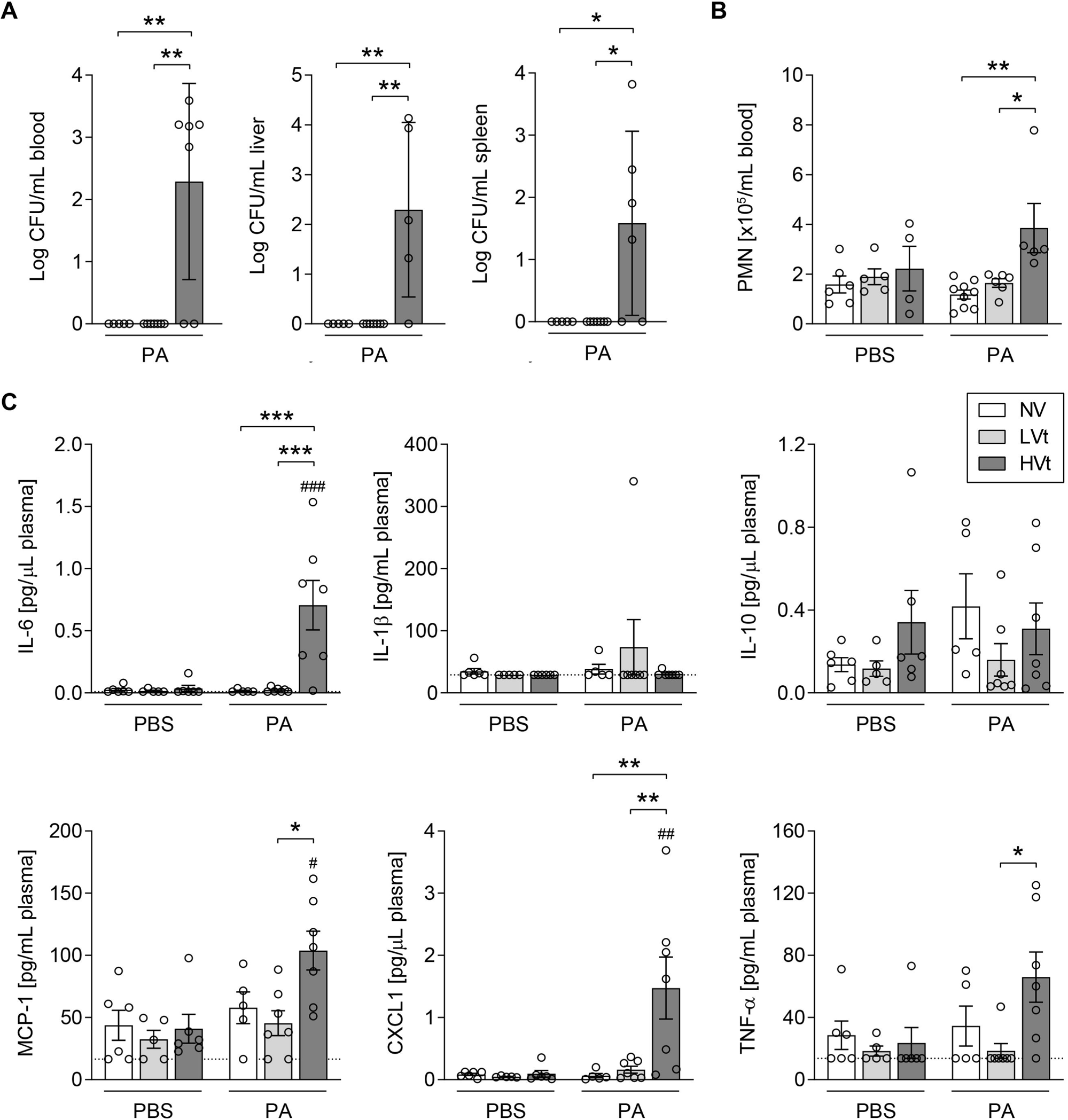
High tidal volume ventilation promotes systemic dissemination of *P. aeruginosa.* Mice were mechanically ventilated with HVt (34 mL/kg), LVt (9 mL/kg) for 4h or ventilated for 10 min (LVt) as NV control. After MV either PA103 or sterile PBS was instilled intratracheally and mice were extubated and left breathing spontaneously for 24h. (**A**) CFUs in blood, liver and spleen homogenates were determined. (**B**) Blood PMNs were differentiated and quantified using a scil Vet abc hematology analyzer. (**C**) Inflammatory cytokines and chemokines in plasma were measured with multiplex bead-based immunoassay technique. Data are presented as mean ± SEM, and analyzed using one-way (**A**) or two-way (**B-C**) ANOVA and Sidak’s multiple comparisons test. In (**A**) and (**C**), n = 5-7 per group; in (**B**), n = 4-6 (PBS) or n = 5-9 (PA). ^#^ indicates significant difference compared with respective sham-infected group, * as indicated. *^/#^*p*<0.05, **^/##^*p*<0.01, ***^/###^*p*<0.001. *CFUs,* colony-forming units; *CXCL*, C-X-C motif chemokine ligand; *HVt,* high tidal volume; *IL*, interleukin; *LVt,* low tidal volume; *MCP*, monocyte chemoattractant protein; *MV,* mechanical ventilation; *NV*, non-ventilated; *PA, Pseudomonas aeruginosa*; *PBS,* phosphate-buffered saline; *PMN*, polymorphonuclear cells; *TNF*, tumor necrosis factor.

### HVt promotes lung injury upon *P. aeruginosa* pneumonia

Histopathological evaluation revealed that HVt led to distinct pulmonary lesions upon PA infection as illustrated by representative H&E-stained whole lung images (**figure 4A**). As compared to LVt and NV mice, HVt mice infected with PA showed the largest affected lung area and most severe lung inflammation infected with PA (**figure 4B**). Semiquantitative histologic scoring (**figure 4C**) revealed that infected HVt mice had overall more severe lung tissue injury with PMN infiltration, hemorrhage, perivascular edema, and alveolar wall damage when compared to other groups (**figure 4D**). In line with permeability analysis, HVt mice also exhibited increased signs of alveolar edema (**supplementary figure E4**).

**Figure 4:**
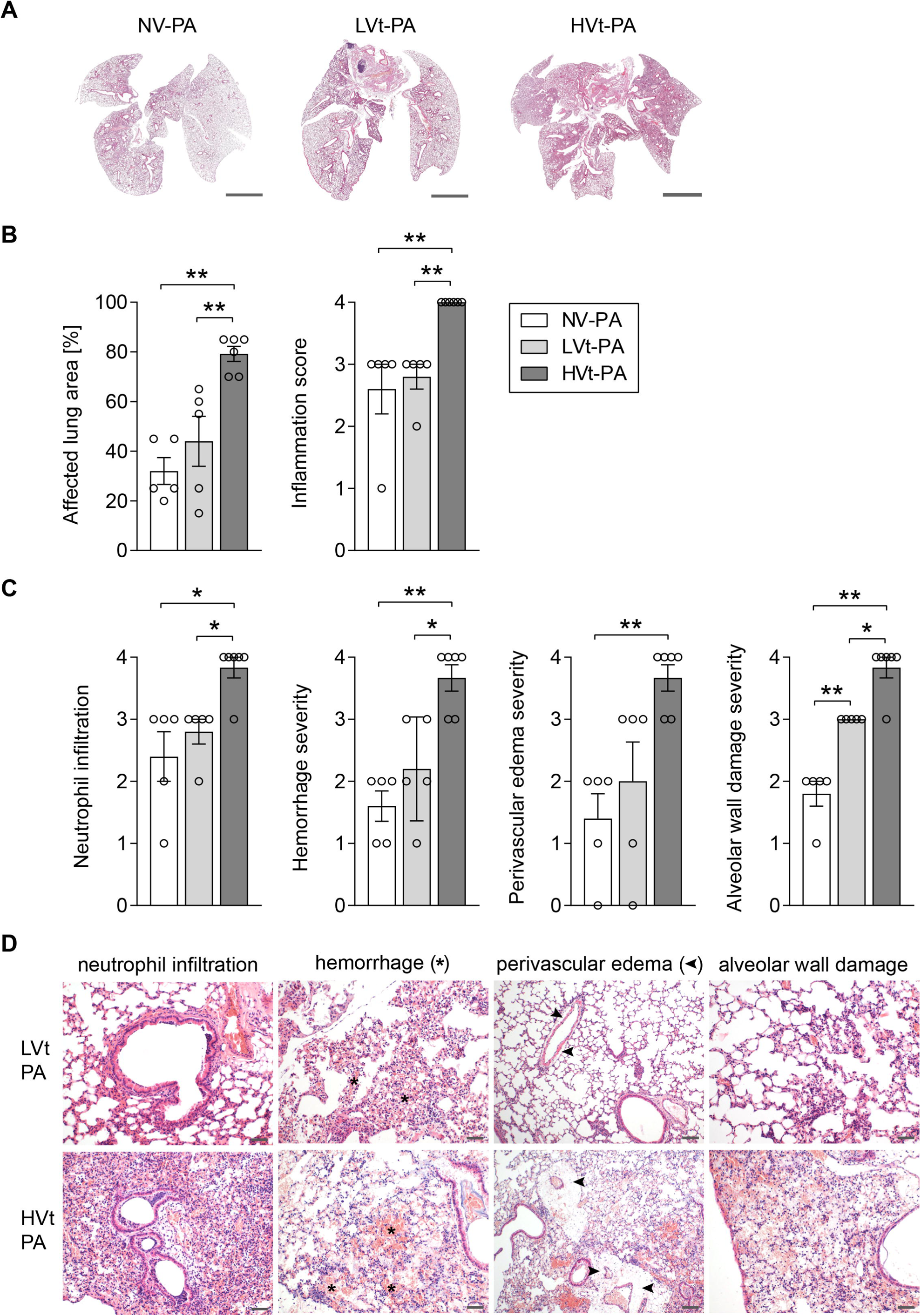
High tidal volume ventilation enhances histological signs of lung injury in subsequent *P. aeruginosa* pneumonia. Mice were mechanically ventilated with HVt (34 mL/kg), LVt (9 mL/kg) for 4h or ventilated for 10 min (LVt) as NV control. After MV either PA103 or sterile PBS was instilled intratracheally and mice were extubated and left breathing spontaneously for 24h. Paraffin-embedded lung sections were stained with hematoxylin and eosin. (**A**) Representative images of whole lung sections from each infected group are shown. Scale bar: 3 mm. (**B**) The percentage of lung area affected by PA pneumonia and total lung inflammation score were calculated. (**C**) Quantification of lung injury: Neutrophil infiltration, hemorrhage, perivascular edema, and alveolar wall damage. All scoring parameters were rated as 0: nonexistent, 1: minimal, 2: mild, 3: moderate, 4: severe. (**D**) Representative images from lung tissue sections of LVt-PA and HVt-PA animals demonstrating histological parameters of lung injury are shown. Scale bar: 50 µm and 100 µm for HVt-PA perivascular edema. In (**B-C**), data are presented as mean ± SEM, n = 5-6 per group. \**p*<0.05, \*\**p*<0.01 (multiple Mann-Whitney U tests with Bonferroni-Holm correction for multiple comparisons). *HVt,* high tidal volume; *LVt,* low tidal volume; *MV,* mechanical ventilation; *NV*, non-ventilated; *PA, Pseudomonas aeruginosa*; *PBS,* phosphate-buffered saline.

### HVt induces intra-alveolar acidification in murine lungs *ex vivo*

To assess the effect of MV on changes in airspace pH as a potential mechanism of increased susceptibility to bacterial infection, we transtracheally sprayed the pH sensitive dye pHrodo and imaged fluorescence in subpleural alveoli of isolated perfused and ventilated lungs (**figure 5A**). Integrated fluorescence intensity of randomly selected alveolar sections was significantly higher in HVt compared to LVt lungs, demonstrating extracellular acidification (**figures 5B, C**).

**Figure 5:**
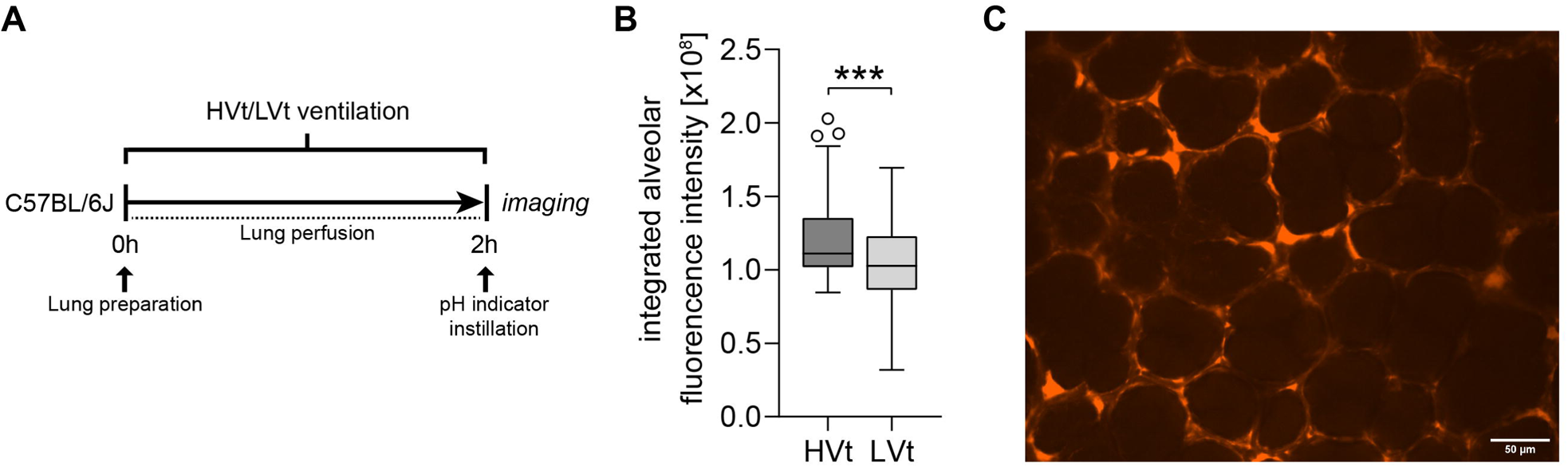
High tidal ventilation induced intra-alveolar acidification in *ex vivo* mice lungs. *Ex vivo* mice lungs were perfused and mechanically ventilated for 2 h with high tidal volume (HVt 20 mL/kg) or low tidal volume (LVt 9 mL/kg). At the end of MV, a pH sensitive indicator with red fluorescence (0.2 µg/µl, Molecular Probes™ pHrodo™ Red Dextran, Thermo Fisher, Berlin, Germany) was applied into lungs using a micro-sprayer and lungs were prepared for microscopy under constant pressure (18 mmHg). (**A**) Experimental design of the *ex vivo* experiments. (**B**) Nine images from 3 independent experiments per group were analyzed. Integrated fluorescence intensity of randomly selected alveolar sections of intermediate size (500-700 µm^2^) was quantified. Data are presented as boxplots depicting median, quartiles and range excluding outliers (open circles), n = 123 (LVt) or n = 150 (HVt). (**C**) Representative image of lung alveoli. *HVt,* high tidal volume; *LVt,* low tidal volume; *MV,* mechanical ventilation.

### Cyclic stretch of lung epithelial cells acidifies epithelial lining fluid and promotes bacterial growth

To test whether HVt-induced acidification of alveolar lining fluid may be due to cyclic stretching of alveolar epithelial cells and contribute to accelerated bacterial growth, we assessed the effect of cyclic stretch on pH, and the role of pH in PA growth on cyclic stretched or non-stretched A549 human lung epithelial cells. Cells were cultured and subjected to 24h cyclic stretch or static conditions, and cells as well as cell-free supernatant (conditioned medium; CM) were collected for infection (**figure 6B**). After stretch, the pH of stretched cell supernatant was significantly more acidic (7.30±0.01) when compared to supernatant of static cells (7.62±0.09) (**figure 6B**). Static or stretched cells were separately seeded onto new culture plates and replenished with CM from either stretched or static conditions. Finally, cells were infected with GFP-PAO1 strain (MOI 10) for 4h. Cells were washed and cells and adhering bacteria were stained for microscopy. Notably, the ratio of bacteria/cell increased only in infected A549 cells that were cultured with CM of stretched, yet not with CM of static cells (**figure 6C**). In contrast, the previous culture conditions of the cells (static vs. stretched) did not impact bacterial growth. The finding that CM from stretched cells promotes bacterial infection and growth was further substantiated by isolation of more PA from cells cultured with CM from stretched as compared to static cells regardless of MOI (**supplementary figure E5**). Analyzing PA growth 24 h p.i. as a function of pH in untreated, acidified, or alkalized medium at pH of 7.41±0.03, 6.92±0.05, and 7.96±0.03, respectively (**figure 6D**), bacterial replication was higher in acidified medium compared to untreated or alkalized medium. Likewise, PA growth was higher in cell-free acidified as compared to untreated or alkalized medium. The finding of stretch-induced acidification and the resulting increase in bacterial growth was reproduced in hpAECs (**supplementary figure E6**). These findings demonstrate that even a relatively modest acidification of the epithelial lining fluid as induced by cyclic stretch of lung epithelial cells can profoundly increase PA growth.

**Figure 6:**
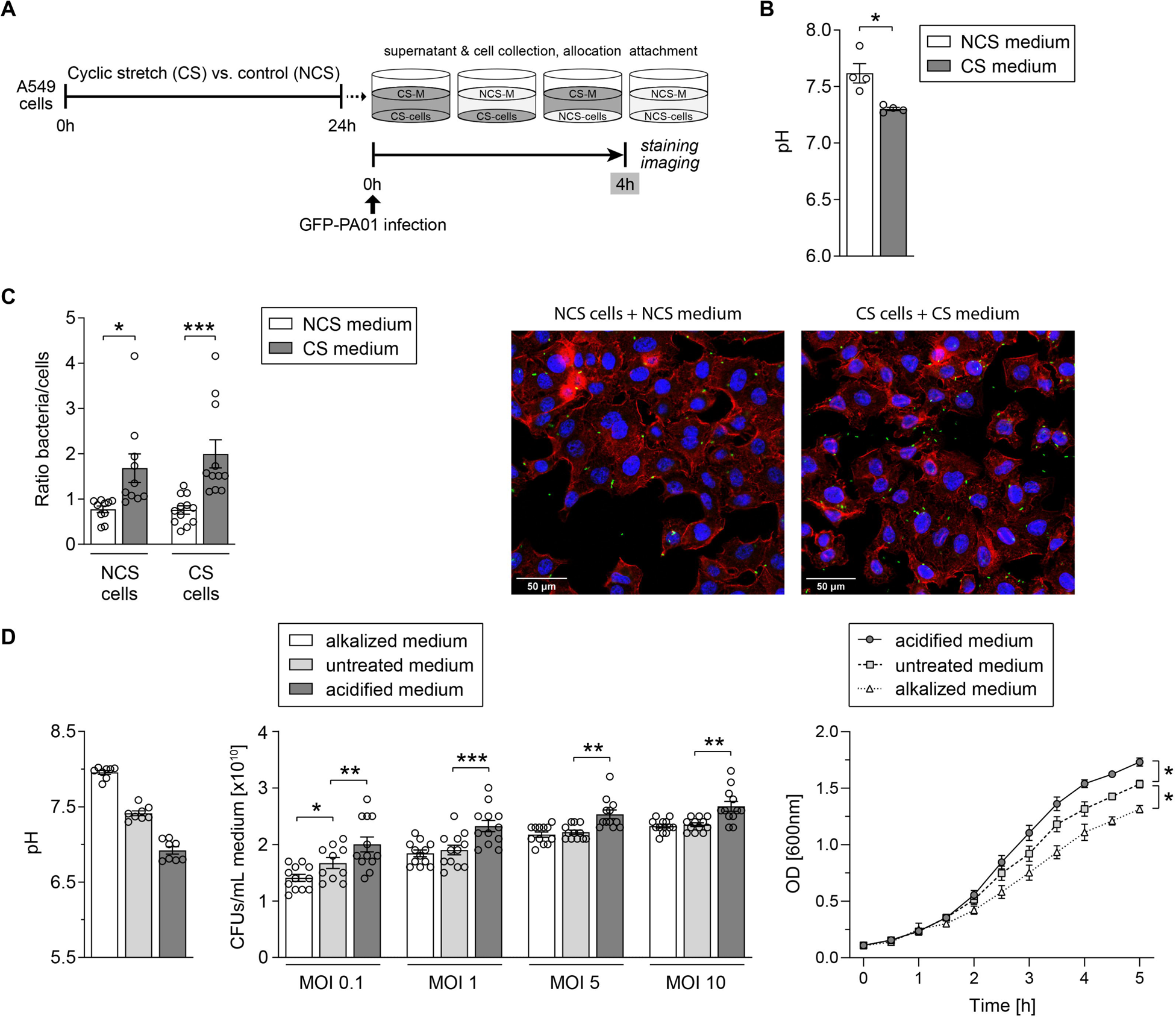
Cyclic stretch of lung epithelial cells acidifies epithelial lining fluid and promotes bacterial adhesion and growth. (**A**) *In vitro* experimental design: cyclically stretched (CS) or non-stretched (NCS) A549 cells were infected with GFP-PAO1 in stretched or non-stretched medium (epithelial lining fluid) for 4h, washed and cells/adhering bacteria were stained with Phalloidin, DAPI and anti-GFP for immunofluorescence. (**B**) pH of stretched and non-stretched medium. (**C**) Left: Evaluation of n = 10×3×3 tile scans and calculation of the bacteria/cell ratio. Right: Representative images of non-stretched cells in non-stretched medium and stretched cells in stretched medium. (**D**) For bacterial growth experiments with GFP-PAO1, HCl and NaOH were used to prepare acidified or alkalized conditioned medium, respectively. Left: pH of treated and untreated media. Middle: A549 cells were infected with PAO1 at MOI 0.1, 1, 5 and 10 in conditioned media and after 24h of incubation CFUs were determined. Right: Growth curves of GFP-PAO1 in RPMI-1640 conditioned media. Data are presented as mean ± SEM, and analyzed using Mann-Whitney U test (**B**), two-way ANOVA and Sidak’s multiple comparisons test (**C, D** middle) or two-way repeated measures ANOVA (**D** right). In (**B**), n = 4 per group; in (**C**) and (**D**) middle, n = 10-12 per group; in (**D**) left, n = 8 per group; in (**D**) right, n = 5-6 per group. \**p*<0.05, \*\**p*<0.01, \*\*\**p*<0.001. *CFU,* bacterial colony-forming unit; *DAPI,* 4′,6-Diamidino-2-phenylindole dihydrochloride; *GFP,* green fluorescence protein; *HCl,* Hydrochloric acid; *MOI,* multiplicity of infection; *NaOH,* sodium hydroxide; *OD*, optical density; *PA, Pseudomonas aeruginosa*.

### Inhibition of NHE1 expression enhances epithelial lining fluid acidification and bacterial growth

To characterize the mechanism underlying the observed *in situ* and *in vitro* acidification of the epithelial lining in response to overventilation or mechanical stretch, respectively, we analyzed gene expression of ion transporter Na^+^/H^+^ exchanger 1 (NHE1) and cystic fibrosis transmembrane conductance regulator (CFTR), both prominently involved in the regulation of extracellular pH. Indeed, gene expression of both CFTR and NHE1 was downregulated in HVt groups (**figure 7A**). To test whether this downregulation might affect the pH of alveolar surface liquid, we pharmacologically inhibited either CFTR or NHE1 in A549 cells and measured pH alterations after 24h in the cell supernatant. Treatment with the NHE inhibitor cariporide evoked significant cell supernatant acidification, while supernatant pH of cells incubated with CFTR-inhibitor 172 was unaltered (**figure 7B**). In line, NHE1 inhibition led to a significant increase in bacterial growth when compared to control and CFTR inhibition (**figure 7B**), which was also observed in hpAECs (**figure 7C**). Taken together these findings suggest a major role of NHE1 downregulation in airspace acidification and enhanced bacterial growth.

**Figure 7:**
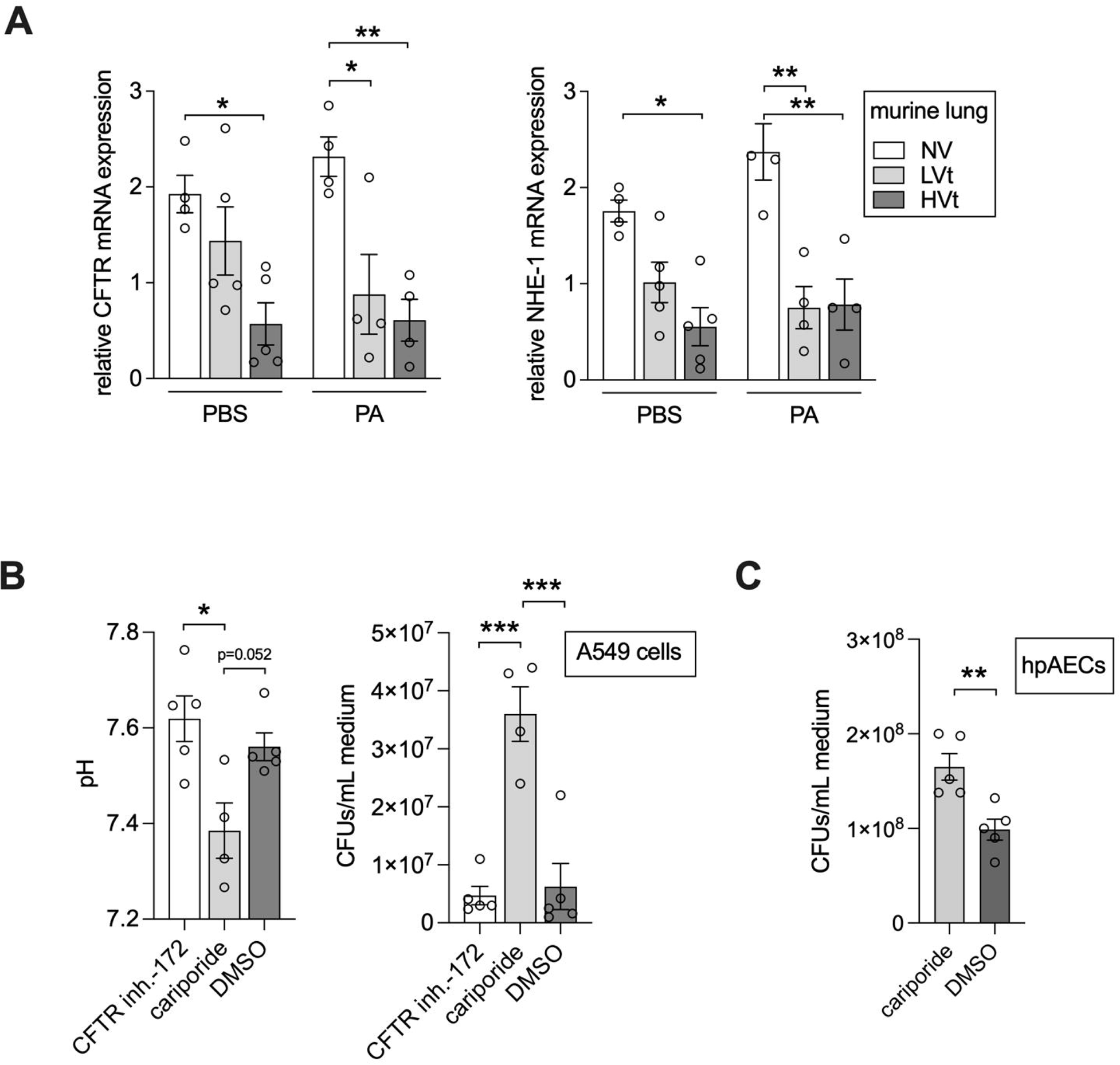
Na+ / H+ exchanger 1 (NHE1) is downregulated upon mechanical ventilation *in vivo* and NHE1 inhibition increased airway liquid acidification promoting bacterial growth *in vitro*. (**A**) Mice were mechanically ventilated with HVt (34 mL/kg), LVt (9 mL/kg) for 4h or ventilated for 10 min (LVt) as NV control. After MV either PA103 or sterile PBS was instilled intratracheally and mice were extubated and left breathing spontaneously for 24h. Relative gene expression of CFTR and NHE1 in lung homogenates was determined and analyzed. (**B**) A549 cells were treated with CFTR inhibitor-172, NHE1 inhibitor cariporide (10 mM) and dimethyl sulfoxide (DMSO) solvent as control. Medium pH was measured after 24h incubation and cells were infected with PAO1 at the MOI of 10 for 4h. CFUs in the medium were determined the next day. (**C**) Bacterial growth experiments were repeated with human primary alveolar epithelial cells (hpAEC). Data are presented as mean ± SEM, and analyzed using two-way (**A**) or one-way (**B**) ANOVA and Sidak’s multiple comparisons test or Mann-Whitney U test (**C**); n = 4-5 per group. \**p*<0.05, \*\**p*<0.01, \*\*\**p*<0.001. *CFTR*, cystic fibrosis transmembrane conductance regulator; *CFUs,* colony-forming units; *DMSO,* dimethyl sulfoxide; *HVt,* high tidal volume; *hpAEC,* human primary alveolar epithelial cells; *inh.*, inhibitor; *LVt,* low tidal volume; *MOI,* multiplicity of infection; *MV,* mechanical ventilation; *NHE1*, Sodium / Hydrogen exchanger 1; *NV*, non-ventilated; *PA, Pseudomonas aeruginosa*; *PBS*, phosphate-buffered saline.

### Pharmacological inhibition of NHE1 increases *P. aeruginosa* pneumonia severity in protectively ventilated mice

To test for the functional relevance of the observed acidification of alveolar lining fluid subsequent to NHE1 inhibition, we treated mice with 3 mg/kg cariporide i.v. before subjecting them to non-injurious MV with low tidal volume (LVt 9 mL/kg) and subsequent infection with PA (**figure 8A**). Compared to other groups, body temperature decreased in mice treated with cariporide and subjected to LVt ventilation (**figure 8B**), while pulmonary bacterial growth was approximately tripled (**figure 8C**). Analysis of inflammatory cells in BALF and lungs revealed no significant differences in PMN and AM counts between the groups. Yet, the concentrations of inflammatory cytokines (MCP-1, TNF-α, GM-CSF, CXCL-5) showed significant elevations in BALF of LVt cariporide treated mice as compared to non-treated LVt and NV mice (**figure 8E**). Likewise, alveolo-capillary permeability significantly increased in LVt and cariporide treated mice as compared to other groups (**figure 8F**), suggesting that cariporide treatment led to an increased inflammatory response and lung barrier disruption upon PA infection. Notably, however, extra-pulmonary dissemination of PA into blood, liver and spleen (**supplementary figure E7A**) and systemic inflammatory response markers (**supplementary figure E7B-C**) were not altered by cariporide treatment, which may indicate that loss of NHE1 function by itself is sufficient to increase lung pneumonia severity, yet without promoting its systemic dissemination in the absence of additional mechanical stressors such as high tidal volume ventilation.

**Figure 8:**
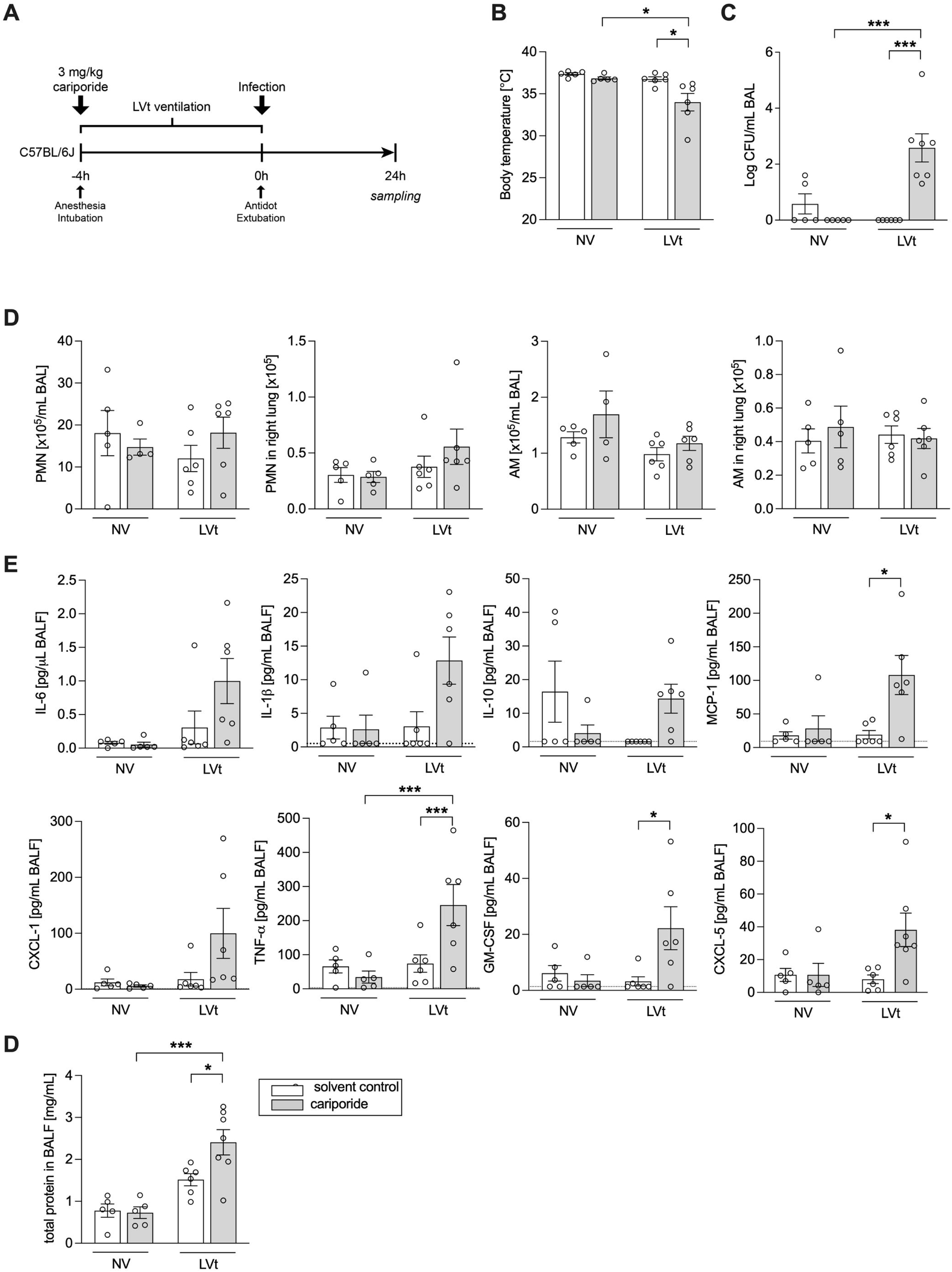
NHE1 inhibition increased susceptibility of mice to *P. aeruginosa* infection after low tidal ventilation. Mice were anesthetized, treated with either 3 mg/kg cariporide or solvent iv and mechanically ventilated with LVt (9 mL/kg) for 4h or ventilated for 10 min (LVt) as NV control. After MV either PA103 or sterile PBS was instilled intratracheally and mice were extubated and left breathing spontaneously for 24h. (**A**) Experimental design of the *in vivo* cariporide treatment experiments. (**B**) Body temperature was measured at the end point of the experiment. (**C**) Pulmonary bacterial colony-forming units (CFUs) in BAL were determined. (**D**) Numbers of PMNs and alveolar macrophages in BAL and lung homogenates were differentiated and quantified *via* flow cytometry. (**E**) Inflammatory cytokines and chemokines in BALF were measured with multiplex beads-based immunoassay technique. (**F**) Alveolar-capillary permeability was assessed by measuring total BAL protein. n = 5-7 per group. Data are presented as mean ± SEM, and analyzed using two-way repeated measures ANOVA and Sidak’s multiple comparisons test. ^#^ indicates significant difference compared with respective sham-infected group, * as indicated. \*\**p*<0.01, \*\*\**p*<0.001. *BAL,* bronchoalveolar lavage; *CXCL*, C-X-C motif chemokine ligand; *IL*, interleukin; *LVt,* low tidal volume; *MV,* mechanical ventilation; *NV*, non-ventilated; *PMN*, polymorphonuclear cells; *TNF*, tumor necrosis factor.

## Discussion

This study demonstrates that high tidal volume ventilation (HVt) promotes bacterial growth and pneumonia severity upon subsequent lung infection by *P. aeruginosa* (PA). Our *ex vivo* and *in vitro* results collectively suggest that cyclic stretch of epithelial cells causes acidification of alveolar lining fluid, probably via NHE1 downregulation, promoting pulmonary PA adhesion and growth, and accelerating the development of pneumonia and systemic bacterial dissemination.

As compared to spontaneously breathing patients, the incidence of nosocomial pneumonia is higher in mechanically ventilated patients. This so-called ventilator-associated pneumonia (VAP) is most commonly induced by PA (4). VAP frequently hampers success in ARDS treatment (23), stressing the need for improved mechanistic understanding to enable the development of novel therapeutic strategies. However, complex underlying clinical entities and ambiguous diagnostic criteria have marred the success of clinical research on VAP development specifically in ARDS patients (24, 25). Therefore, various animal models have been established to study VAP. While in some studies, mice were infected prior to or at the onset of MV (15, 16, 18), our present model comprises antagonization of anesthesia, recovery of consciousness and extubation after mechanical ventilation, and infection with PA followed by spontaneous breathing for 24h. As such, the model presented here mimics the characteristic sequence of events in VAP, i.e. first MV, then infection. This clinical course of human VAP has previously been reproduced exclusively in a rat and a porcine model (19) to which our newly established murine model might serve as a versatile alternative, particularly in view of opportunities for mechanistic studies by use of transgenic mice. Further, as a consequence of HVt, PA bacteremia and sepsis-related extra-pulmonary organ damage developed, which are often associated with treatment failure in PA-VAP patients (26). Moreover, we observed model-specific consistencies and differences regarding the effects of ventilation. Bacterial burden was severely increased upon HVt in our present model, comparable to previous reports in rat and swine (18, 27). On the other hand, LVt has previously been reported to induce inflammatory responses in blood and lungs of rats (18). Protective ventilation of healthy lungs in our present mouse model did not evoke measurable injury after 24h, which underscores its translational relevance and hence, its suitability to investigate mechanisms of VAP pathogenesis in greater detail.

In this study, tidal volume was set at 34 mL/ per kg bodyweight for the HVt protocol. According to the baby lung concept, even protective ventilation strategies may result in highly increased stress and strain in overinflated regions of ARDS lungs. This pathological scenario is commonly mimicked in experimental research by subjecting laboratory animals to short-term high tidal-volume ventilation (21, 28).

Innate immune activation is critical for defending PA infection in mechanically ventilated patients (29) and animal experiments (30). Any impairments of host innate immune defense, particularly of antimicrobial functions, due to airspace acidification are likely to favor pulmonary PA colonization and growth (11, 31, 32). Clinical studies analyzing exhaled breath condensates showed that decreased pH levels in patients with chronic inflammatory airway diseases corresponded to increased PA growth and induced sputum neutrophil counts (33). Moreover, Torres and coworkers demonstrated that deliberately acidified infection doses of PA increased disease severity and correlated with an exaggerated IL-1β-dependent inflammatory response including increased neutrophil recruitment and MPO release (34). However, whether the airway acidification itself, also detected in patients with acute lung injury (ALI) and acute respiratory distress syndrome (ARDS) (7, 8), may facilitate bacterial adherence and the subsequent development of PA pneumonia or is rather a sign for aggravation of pulmonary inflammation remains unclear. Notably, the potential tripartite relationship between cyclic stretch of airway epithelial cells, pH reduction and bacterial growth advantage has been consolidated *in vitro* (35), yet in vivo evidence for a role of VILI-induced acidification in promoting PA replication and increasing VAP severity remains to be demonstrated.

In the present study, cyclic stretch of lung epithelial cells caused acidification of epithelial lining fluid and augmented bacterial adhesion and growth. Moreover, the pH in lungs from mice subjected to HVt was found to be acidic compared to spontaneously breathing mice. The acidic milieu in the airspace is expected to confer a significant growth advantage for PA under the conditions tested and might in parallel impact the physiology of PA as several studies indicate that adaptation to acidic pH could increase virulence and induce drug/antibiotic tolerance in PA (36, 37). The chloride channel CFTR and the sodium-proton exchanger NHE1 are expressed in airspace epithelial cells (38, 39) and involved in extracellular pH regulation (12). While a lack of CFTR may lead to airway surface liquid acidification and higher likelihood for PA infections (11), the impact of NHE1 on pH regulation and pulmonary infections was so far unknown. In our murine model, we observed substantial downregulation of CFTR and NHE1 expression in lung tissue after MV and infection. Interestingly, inhibition of NHE1 but not CFTR evoked acidification and subsequently enhanced bacterial growth on human alveolar epithelial cells, providing a possible underlying mechanism for the observed airspace acidification and increased susceptibility to PA *in vivo*. Notably, in a very recent study acidity increased biofilm formation and antibiotic resistance of PA, which was reversed by reestablishing pH-neutral conditions (40). Further corroborating these results, *in vivo* inhibition of NHE1 with cariporide significantly increased bacterial burden and disease severity in PA infections in LVT ventilated animals compared to solvent treated controls, while the systemic dissemination and inflammatory cell recruitment were not affected. Notably, this could be indicative that additional effects of MV on lung lung barrier ultrastructure, may be required for the systemic spread of the infection (**supplementary figure E8**). These results further underpin that NHE1-regulated airspace acidification has a significant effect on pulmonary growth of Pseudomonas aeruginosa.

In our study, increased bacterial growth was associated with enhanced induction of pro-inflammatory cytokines (e.g., IL-1ß, CXCL-1, IL-6 and MCP-1) and neutrophil activation as well as pronounced lung inflammation and injury in HVt-PA mice, reflecting increased disease severity.

This study has several limitations. First, it is noteworthy that the observed acidification upon cell stretch is moderate when compared to the artificially conditioned media for bacterial growth experiments. Nonetheless, measurements were performed in total supernatant, suggesting that actual changes in the thin layer of pulmonary epithelial lining in situ may possibly be more drastic. Additionally, increased stretch-induced cell death as possible cause for acidification was excluded (**supplementary figure S9**). Second, unlike the short duration of ventilation used in the present model, VAP patients are continuously ventilated over the progression of pneumonia. Therefore, long-term effects of HVt and associated VAP on immune functions, disease progression, as well as morbidity and mortality cannot be directly inferred from our mouse model. That notwithstanding, the present model replicates key clinical features of PA-VAP reaffirming its utility as a model to investigate the complex mechanisms of MV-induced PA susceptibility and VAP onset.

In conclusion, employing our novel murine model of PA-VAP we were able to show that high tidal volume ventilation increases the susceptibility of mice to pursue pseudomonas aeruginosa pneumonia. We further provide evidence that cyclic cell stretch and high tidal volume ventilation result in airspace acidification paving the way to pneumonia upon PA infection. Ultimately, we identified NHE1 downregulation as critical mechanism of alveolar acidification and possible contributor to VAP pathogenesis. Gaining further insights into the processes of acidification and VAP development may identify novel therapeutic targets to combat this potentially lethal complication in intensive care medicine.

## Acknowledgments

We thank Ulrike Behrendt Katharina Hellwig and Denise Bartel for their excellent technical assistance.

## Authors’ contributions

MF and CT performed the animal and bacterial culture experiments, analyzed the data and drafted the manuscript. MF and MW planned and supervised the study, interpreted the data and revised the manuscript. JiL performed bacterial culture experiments and revised the manuscript. AH and SF performed *in vitro* microscopy experiments and revised the manuscript. WMK and QZ performed *ex vivo* lung microscopy experiments and revised the manuscript. CB and MO performed electron microscopy experiments and revised the manuscript. GK, EL and HCMR interpreted the data and edited the manuscript. JaL performed statistical analysis and revised the manuscript. ADG and TCB performed histological examination and revised the manuscript. All authors read and approved the final manuscript version.

None of the authors has a financial relationship with a commercial entity that has an interest in the subject of this manuscript.

## Declaration of interests

The authors declare no competing interests.

## Supplementary figure legends

**Supplementary figure E1:**
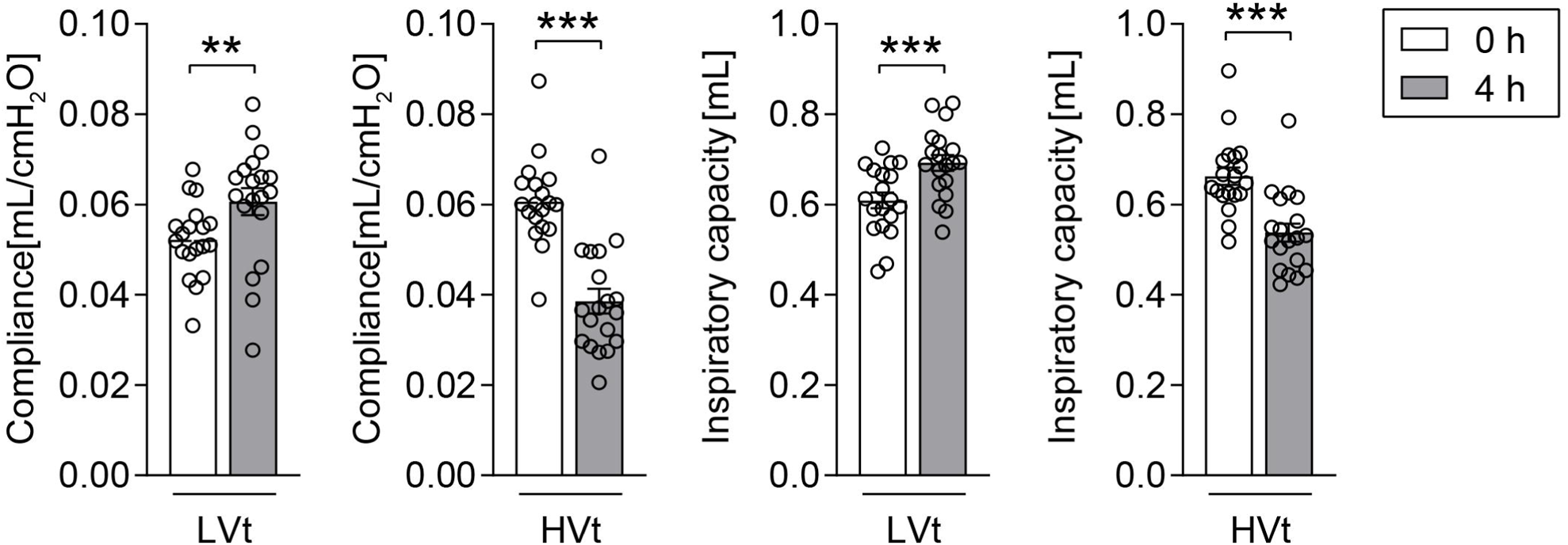
High tidal volume ventilation induces ventilator-induced lung injury in the murine model of HVt ventilation. Mice were mechanically ventilated for 4h with high tidal volume (HVt 34 mL/kg) or low tidal volume (LVt 9 mL/kg). Static compliance and inspiratory capacity were measured and recorded *via* recruitment maneuvers at the beginning (0 h) and end of MV (4h). Data are presented as mean ± SEM; n=19 per group. \*\**p*<0.01, \*\*\**p*<0.001 (Wilcoxon signed ranked test). *HVt,* high tidal volume; *LVt,* low tidal; *MV,* mechanical ventilation.

**Supplementary figure E2:**
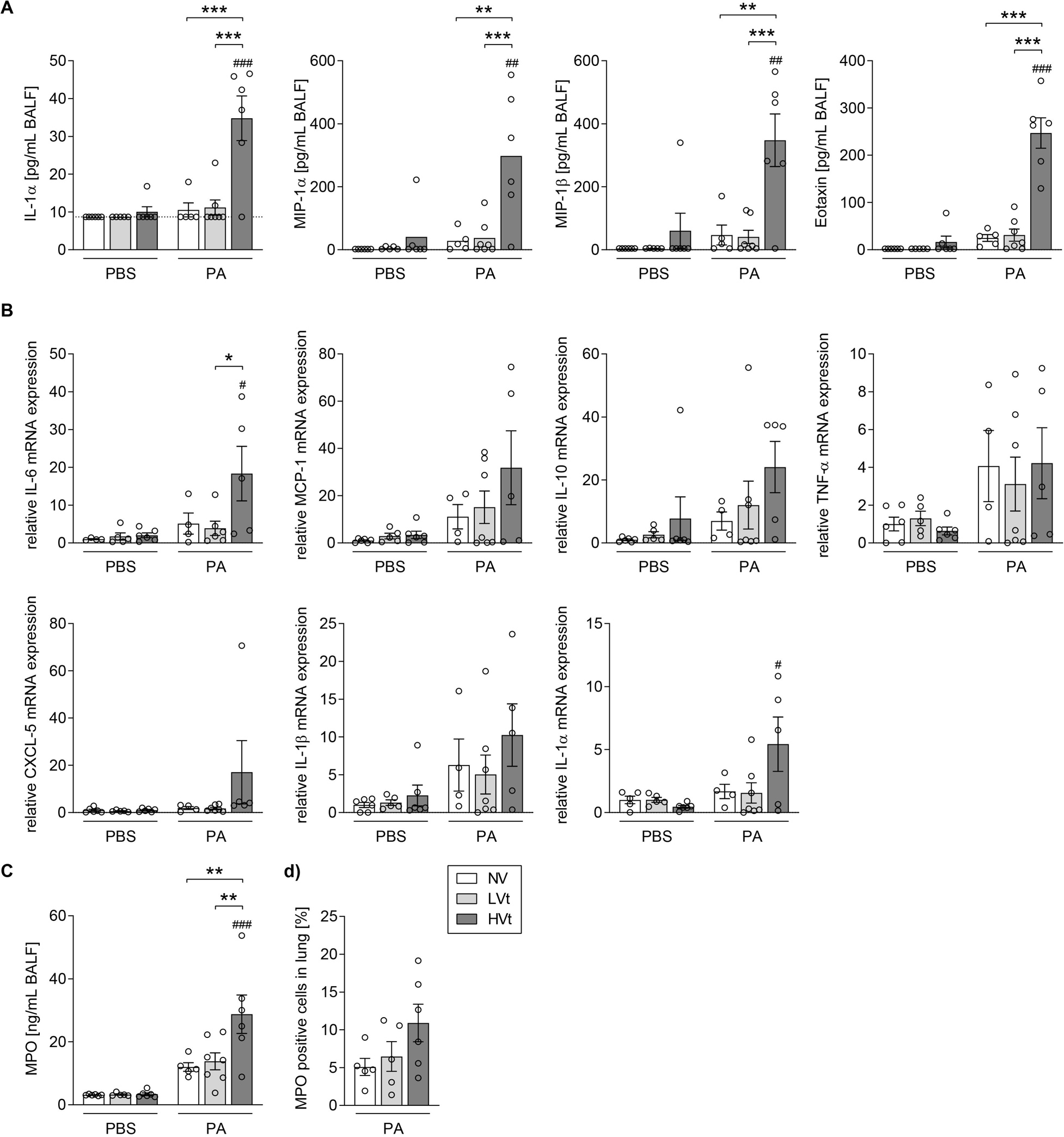
Pulmonary inflammation is increased in *P. aeruginosa* pneumonia subsequent to high tidal volume ventilation. Mice were mechanically ventilated with HVt (34 mL/kg), LVt (9 mL/kg) for 4h or ventilated for 10 min (LVt) as NV control. After MV either PA103 or sterile PBS was instilled intratracheally and mice were extubated and left breathing spontaneously for 24h. (**A**) Inflammatory cytokines and chemokines in BALF were measured with multiplex bead-based immunoassay technique. (**B**) Relative gene expression of inflammatory cytokines and chemokines from lung homogenates was determined and analyzed. N = 4-6 (PBS) or n = 4-7 (PA). Data are presented as mean ± SEM. ^#^ indicates significant difference compared with respective sham-infected group, * as indicated. *^/#^*p*<0.05, **^/##^*p*<0.01, ***^/###^*p*<0.001 (two-way ANOVA and Sidak’s multiple comparisons test). *BALF,* bronchoalveolar lavage; *CXCL*, C-X-C motif chemokine ligand; *HVt,* high tidal volume; *IL*, interleukin; *LVt,* low tidal volume; *MCP*, monocyte chemoattractant protein; *MIP*, Macrophage Inflammatory Protein; *MV,* mechanical ventilation; *NV*, non-ventilated; *PA, Pseudomonas aeruginosa*; *PBS,* phosphate-buffered saline; *TNF*, tumor necrosis factor.

**Supplementary figure E3:**
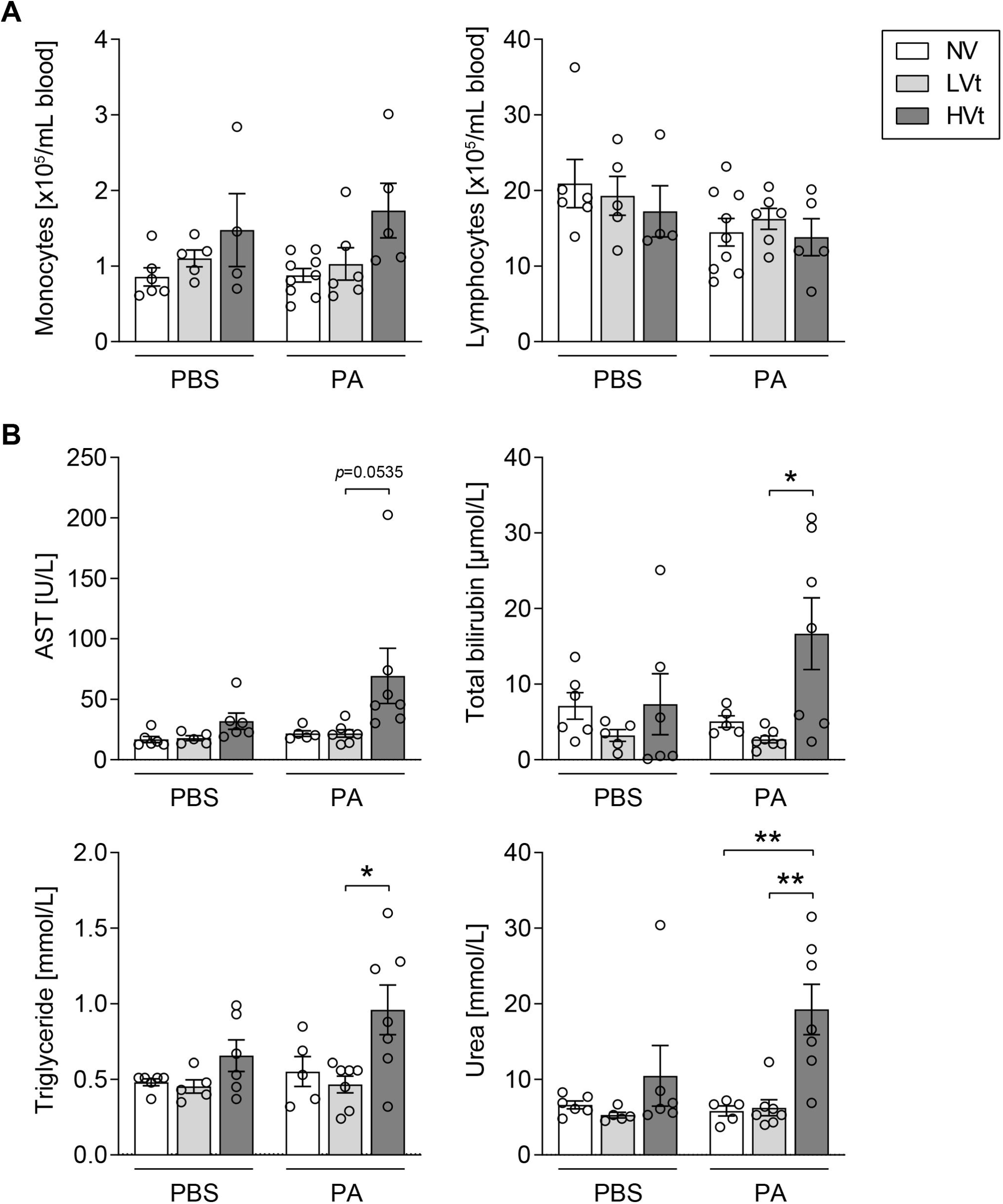
High tidal volume ventilation promotes systemic spread of *P. aeruginosa*. Mice were mechanically ventilated with HVt (34 mL/kg), LVt (9 mL/kg) for 4h or ventilated for 10 min (LVt) as NV control. After MV either PA103 or sterile PBS was instilled intratracheally and mice were extubated and left breathing spontaneously for 24h. Plasma levels of AST, total bilirubin, triglyceride, and urea were determined to evaluate liver and kidney function. n = 5-6 (PBS) or n = 5-7 (PA). Data are presented as mean ± SEM. \**p*<0.05, \*\**p*<0.01 (two-way ANOVA and Sidak’s multiple comparisons test). *AST*, aspartate transaminase, *HVt,* high tidal volume; *LVt,* low tidal volume; *MV,* mechanical ventilation; *NV*, non-ventilated; *PA, Pseudomonas aeruginosa*; *PBS,* phosphate-buffered saline.

**Supplementary figure E4:**
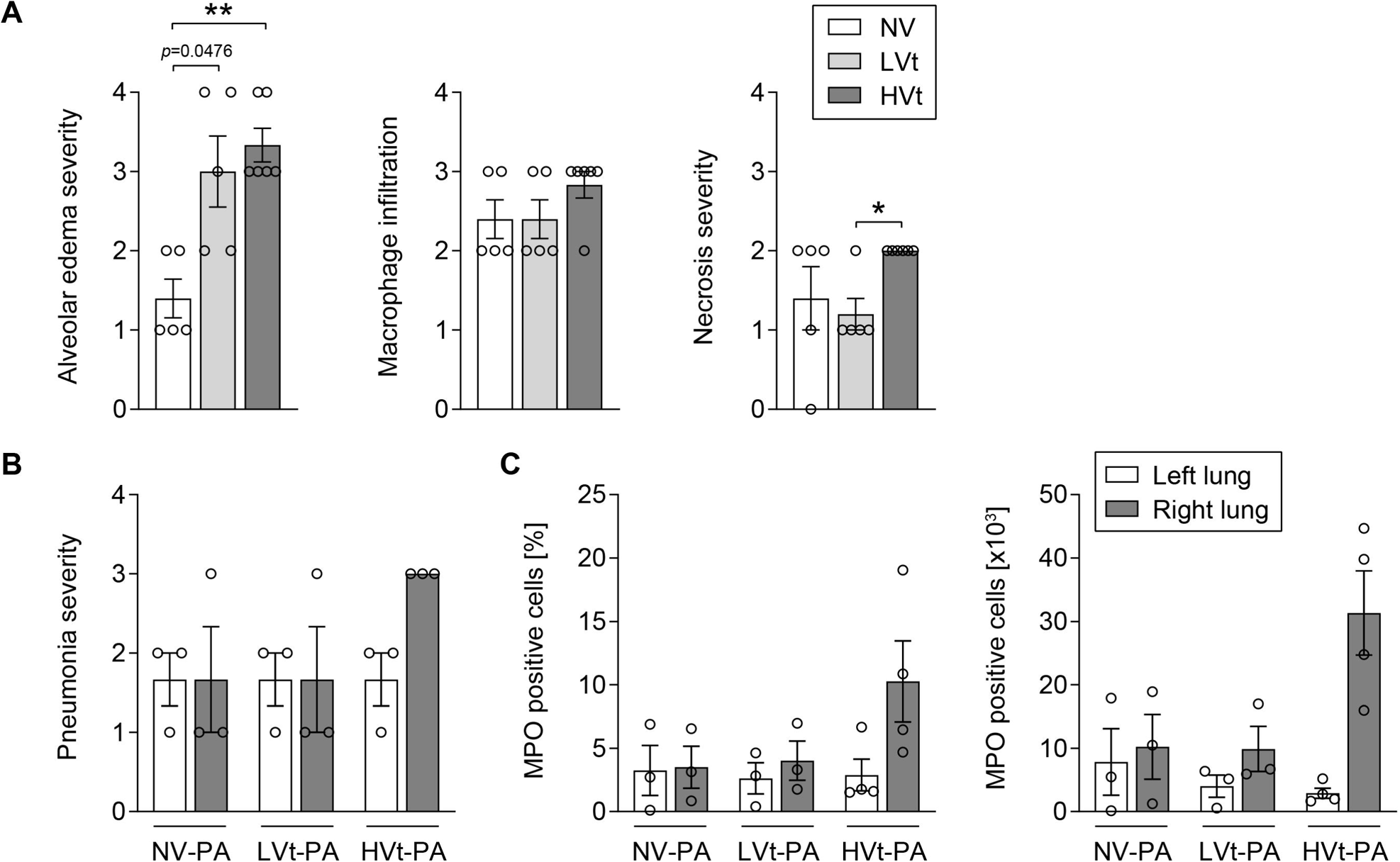
High tidal volume ventilation enhances histological signs of lung injury in subsequent *P. aeruginosa* pneumonia. Mice were mechanically ventilated with HVt (34 mL/kg), LVt (9 mL/kg) for 4h or ventilated for 10 min (LVt) as NV control. After MV either PA103 or sterile PBS was instilled intratracheally and mice were extubated and left breathing spontaneously for 24h. Paraffin-embedded lung sections were stained with hematoxylin and eosin. Alveolar edema, macrophage infiltration and necrosis were evaluated to quantify lung injury. All scoring parameters were rated as 0: nonexistent, 1: minimal, 2: mild, 3: moderate, 4: severe. n = 5-6 per group. Data are presented as mean ± SEM, and analyzed using multiple Mann-Whitney U tests with Bonferroni-Holm correction for multiple comparisons. \**p*<0.05, \*\**p*<0.01. *HVt,* high tidal volume; *LVt,* low tidal volume; *MV,* mechanical ventilation; *NV*, non-ventilated; *PA, Pseudomonas aeruginosa*; *PBS,* phosphate-buffered saline.

**Supplementary figure E5:**
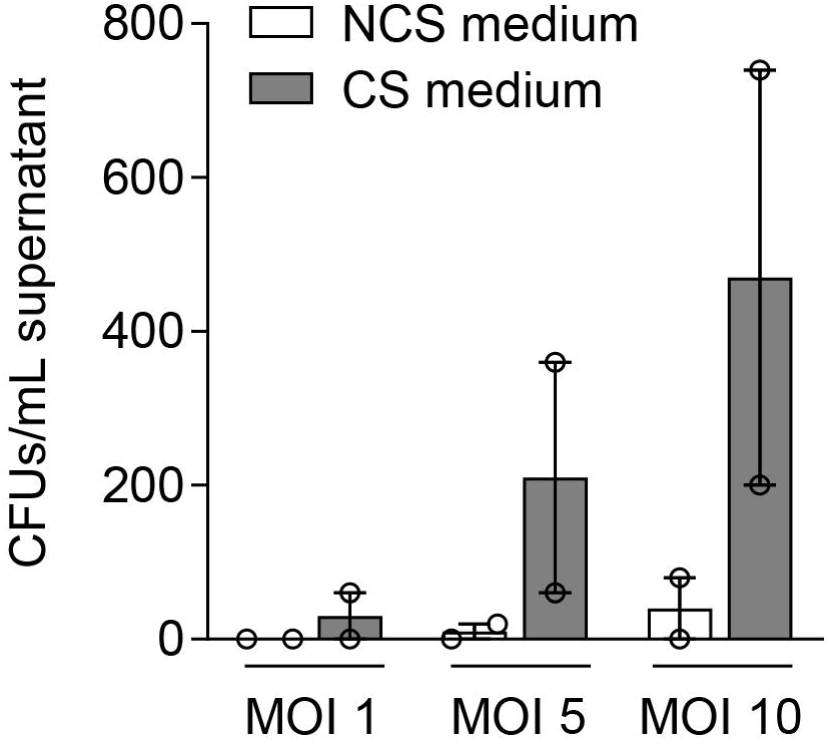
Cyclic stretch of lung epithelial cells promotes bacterial adhesion and growth. Medium from cyclically stretched (CS) or non-stretched (NCS) A549 (experimental scheme figure 6A) was collected. A549 cells were infected with PAO1 at MOI 1, 5 and 10 in respective media and after 24h of incubation CFUs were determined. Data are presented as mean ± SEM of 2 independent experiments. *CFUs,* colony-forming units; *MOI,* multiplicity of infection.

**Supplementary figure E6:**
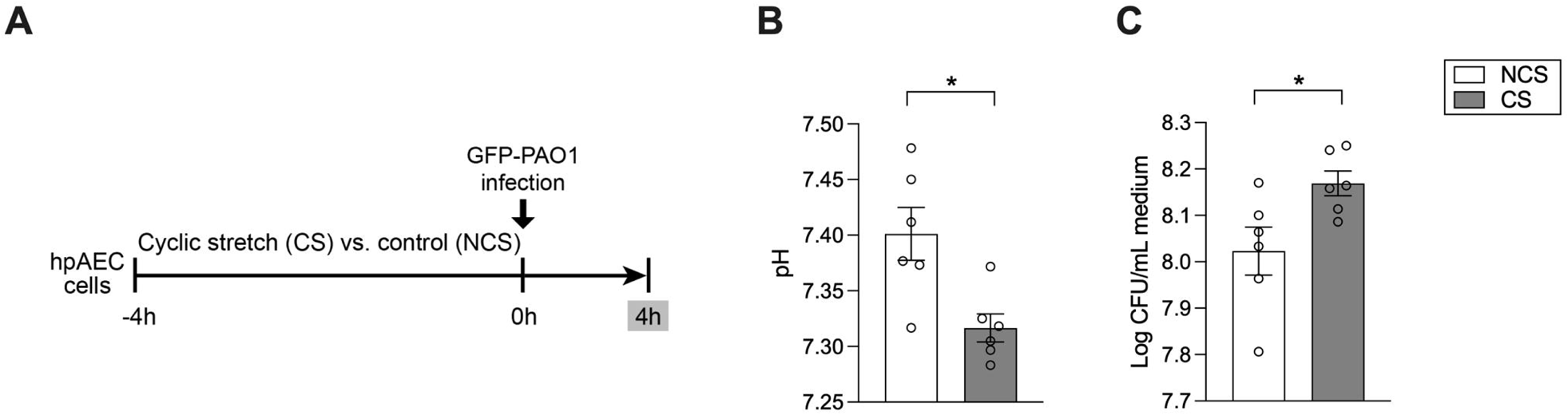
Cyclic stretch of human primary lung alveolar epithelial cells acidifies epithelial lining fluid and promotes bacterial adhesion and growth. (**A**) *In vitro* experimental design: cyclically stretched (CS) or non-stretched (NCS) human alveolar epithelial cells (hpAEC) were infected with GFP-PAO1 and after 4h of incubation CFUs were determined. (**B**) pH of stretched and non-stretched medium. (**C**) CFUs in medium after 24h. Data are presented as mean ± SEM, and analyzed using Mann-Whitney U test, n = 6 per group. \**p*<0.05. *CFU,* bacterial colony-forming unit; *GFP,* green fluorescence protein; *PA, Pseudomonas aeruginosa*.

**Supplementary figure E7:**
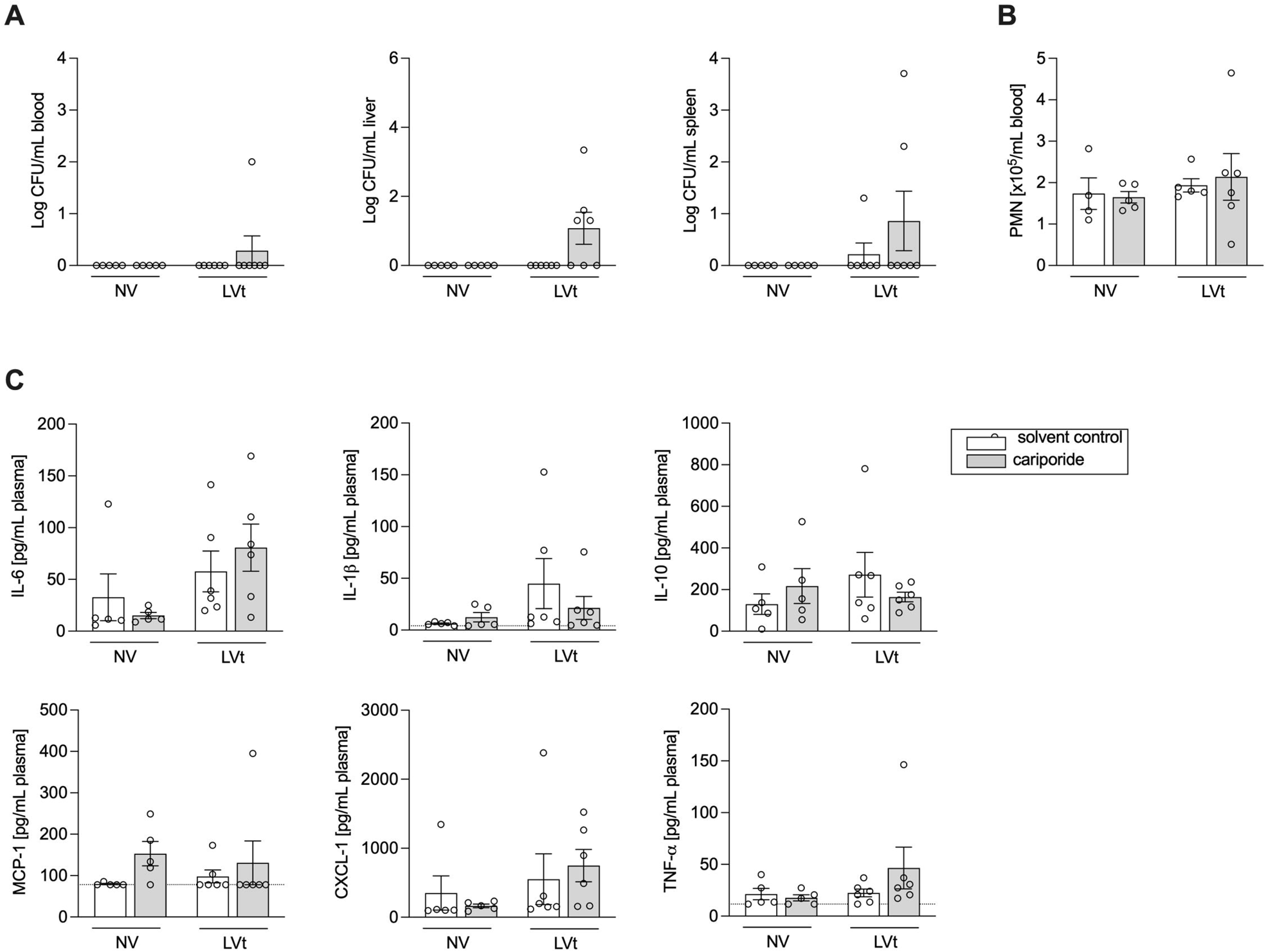
NHE1 inhibition did not promote systemic dissemination of *P. aeruginosa*. Mice were anesthetized, treated with either 3 mg/kg cariporide or solvent iv and mechanically ventilated with LVt (9 mL/kg) for 4h or ventilated for 10 min (LVt) as NV control. After MV either PA103 was instilled intratracheally and mice were extubated and left breathing spontaneously for 24h. (**A**) CFUs in blood, liver and spleen homogenates were determined. (**B**) Blood PMNs were differentiated and quantified using a scil Vet abc hematology analyzer. (**C**) Inflammatory cytokines and chemokines in plasma were measured with multiplex bead-based immunoassay technique. Data are presented as mean ± SEM and analyzed using two-way ANOVA and Sidak’s multiple comparisons test. n = 5-7. *CFUs,* colony-forming units; *CXCL*, C-X-C motif chemokine ligand; *IL*, interleukin; *LVt,* low tidal volume; *MCP*, monocyte chemoattractant protein; *MV,* mechanical ventilation; *NV*, non-ventilated; *PA, Pseudomonas aeruginosa*; *PMN*, polymorphonuclear cells; *TNF*, tumor necrosis factor.

**Supplementary figure E8:**
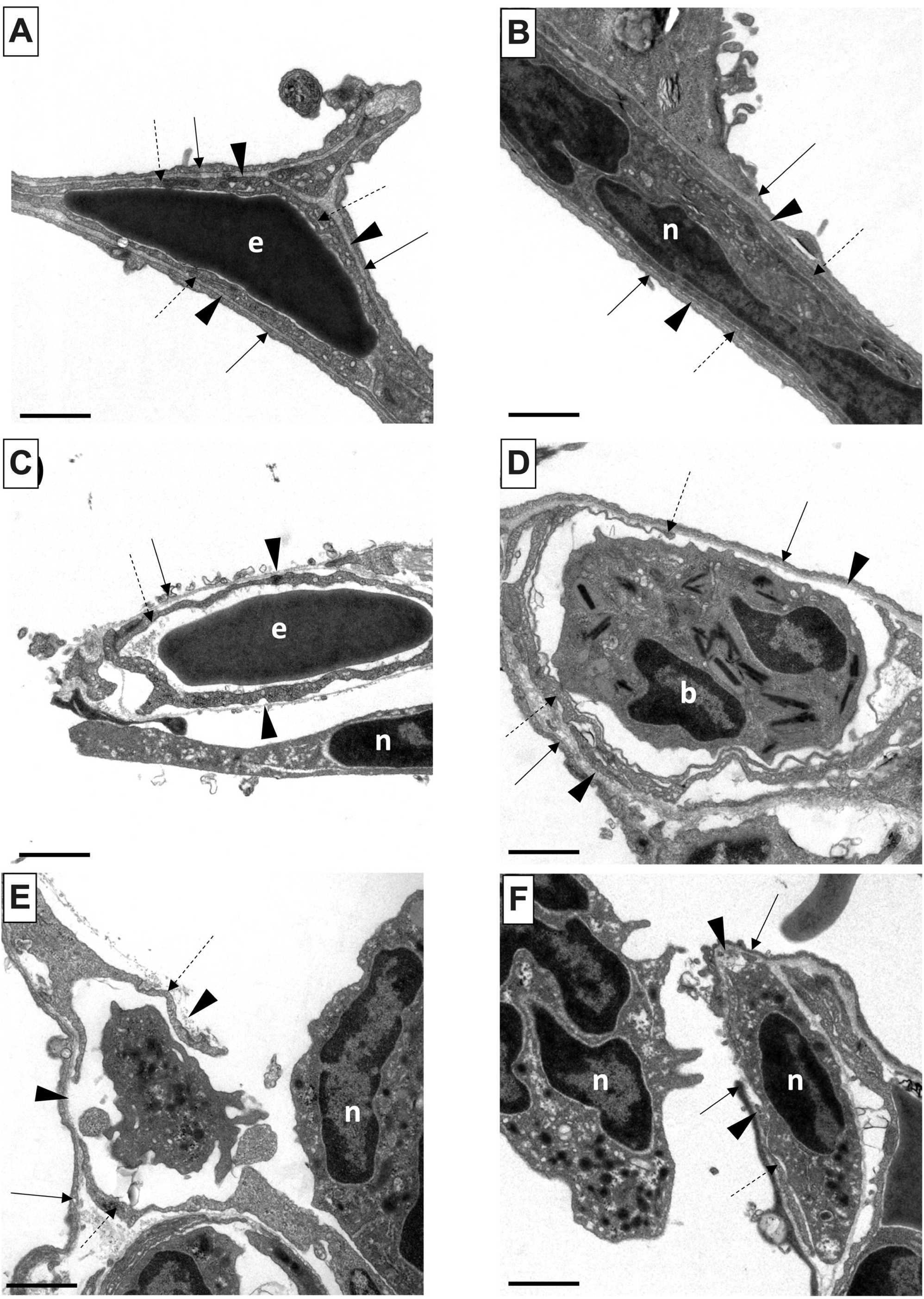
Basement membrane of the air-blood barrier. Mice were mechanically ventilated with HVt (34 mL/kg), LVt (9 mL/kg) for 4h. After MV either PA103 was instilled intratracheally and mice were extubated and left breathing spontaneously for 24h. Lung samples were embedded for electron microscopy. 60-70 randomly acquired images of the air-blood barrier were analyzed per lung. No apparent breaks/discontinuities were observed in the basement membrane in areas with an intact air-blood barrier (**A**, **B**). However, injury to the epithelial (**C**) or endothelial (**D**) cells with partially intact basement membrane or complete distortion of the air-blood barrier (**E**, **F**) was observed in areas with severe injury, which appeared more pronounced in HVT ventilated animals. n, neutrophil; b, basophil; e, erythrocyte; arrow head, basement membrane; arrow, epithelium; dashed arrow, endothelium; scale bars = 1 ρm.

**Supplementary figure E9:**
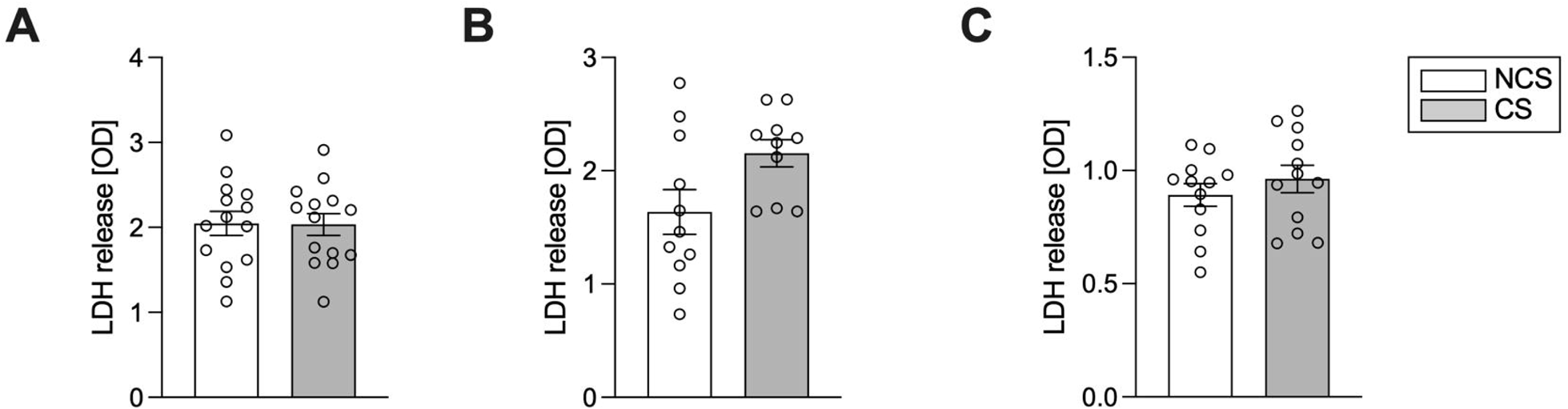
Twenty-four hours cyclic stretch does not induce increased cell death in lung epithelial cells. (**A**) A549 cells, (**B**) human alveolar epithelial cells (hpAEC) and (**C**) H441 lung epithelial cells were exposed to cyclically stretch (CS) or no stretch (NCS) for 24h and LDH concentration as surrogate for cell death was determined via ELISA. Data are presented as mean ± SEM, and analyzed using Mann-Whitney U test, n = 10-14 per group. *LDH*, lactate dehydrogenase.

## Notes

### Competing Interest Statement

The authors have declared no competing interest.

