## Supplementary Materials and Methods for "Overventilation-induced airspace acidification increases susceptibility to Pseudomonas pneumonia"

### Mouse model for ventilator-associated pneumonia (VAP)

All animal experiments were reviewed and approved by the State Office for Health and Social Services, Berlin, Germany. Experimental procedures were carried out in accordance with the European Directive 2010/63/EU on care, welfare and treatment of animals.

Female C57BL/6J mice (8-10 weeks, 18-20 g, Charles River, Sulzfeld, Germany) were anaesthetized (75 µg/kg fentanyl, 1.5 mg/kg midazolam, and 0.75 mg/kg medetomidine; i.p.), orotracheally intubated and mechanical ventilated (FlexiVent; SCIREQ, Montreal, Canada) for 4h with either high tidal volume (HVt: 34 ml/kg) or low tidal volume (LVt: 9 ml/kg) as prior reported (1). Non-ventilated (NV) mice were briefly attached to the ventilator for 10 min in LVt settings to record baseline experimental parameters and thereby served as a control. For NHE1 inhibition experiments 3mg cariporide (Sigma-Aldrich, St. Louis, USA) per kg bodyweight was intravenously injected into the tail vein.

Following an initial recruitment maneuver, the lung function parameters such as mean airway pressure and dynamic elastance were measured and recorded at 5 min intervals during MV by the forced oscillation technique. The terminal recruitment maneuver was performed at 5 min prior the termination of MV. Static compliance and inspiratory capacity were also recorded.

For *Pseudomonas aeruginosa* mice infection studies, 20 µl suspension containing $5 \times 10^4$ colony-forming units (CFU) of *P. aeruginosa* PA103 (American Type Culture

Collection (ATCC) catalog #29260) or 20 µl of sterile phosphate-buffered saline (PBS; Gibco; Thermo Fisher Scientific, Waltham, USA) was instilled via the endotracheal tube.

For post-anesthesia recovery, mice were administered with 0.5 mg/kg flumazenil and 5 mg/kg atipamezole (2), followed by extubation and spontaneous breathing for 24h.

**Bacterial load enumeration and immune cells isolation**

After 24h of infection, anesthetized animals (after administering 80 mg/kg ketamine and 25 mg/kg xylazine i.p.) underwent euthanasia through withdrawal of blood from the *vena cava*. Bronchoalveolar lavage (BAL) was obtained by instilling 800 µl of PBS and draining immediately thereafter. Lungs were flushed with chilled saline (B. Braun, Melsungen, Germany) and left lung was harvested and stored at -80°C for mRNA and protein quantification. Right lung lobes were used for bacterial load enumeration and cell isolation. Lung tissue, BAL fluid (BALF) and blood plasma were stored at -80°C until further analysis. Bacterial CFU counts of BAL, blood and lung, liver and spleen homogenates were enumerated on blood agar after overnight incubation at 37 °C and 5% $CO_2$.

**Blood leukocyte analysis and liver/kidney function tests**

Blood leukocytes were differentiated and quantified by scil Vet abc hematology analyzer (Scil Animal Care Company GmbH, Viernheim, Germany). Plasma levels of liver and kidney function surrogate markers (aspartate aminotransferase, total bilirubin, triglyceride and urea) were analyzed in a clinical laboratory (Laboklin, Melsungen, Germany).

**Lung permeability**

Albumin concentration in BALF and plasma was determined using ELISA (Bethyl Laboratories, Montgomery, USA). Albumin BALF/plasma ratio was used to evaluate alveolar - capillary permeability as previously described (3).

**Flow cytometry analysis**

Flow cytometry was used to identify and individually count BAL and lung leukocytes as previously described (4, 5). Briefly, half right lung was digested in RPMI-1640 medium (Gibco; Thermo Fisher Scientific, Waltham, USA) containing DNAse (AppliChem GmbH, Darmstadt, Germany) and collagenase (Biochrom GmbH; Berlin, Germany). Digested lung lobes were homogenized into a single cell suspension. After blocking the nonspecific binding (anti-CD16/32; Becton, Dickinson and Company, Franklin Lakes, USA), cells were stained with anti-CD11c-APC (N418, Becton, Dickinson and Company, Franklin Lakes, USA), anti-CD11b-PE-Cy7 (M1/70, eBioscience; Thermo Fisher Scientific, Waltham, USA), anti-F4/80-PE (BM8, eBioscience; Thermo Fisher Scientific, Waltham, USA), anti-CD45-FITC (30-F11, Becton, Dickinson and Company, Franklin Lakes, USA), anti-Ly6G-PerCP Cy5.5 (1A8, Becton, Dickinson and Company, Franklin Lakes, USA), anti-Ly6C-BV510 (HK1.4, BioLegend, San Diego, USA), anti-MHCII-AF700 (M6/114.15.2, eBioscience; Thermo Fisher Scientific, Waltham, USA) and anti-SiglecF-BV421 (1RNM44N, eBioscience; Thermo Fisher Scientific, Waltham, USA). Counting beads (CountBright beads, Thermo Fisher Scientific, Waltham, USA) were used to quantify leukocytes according to manufacturer's instructions. Cells were sorted by FACS Canto II flow cytometer (BD Bioscience; USA). Data were analyzed with FlowJo software (FlowJo LLC, Ashland, USA).

**Cytokine and chemokine quantification**

Concentrations of inflammatory cytokines (Interleukin-6 (IL-6), Interleukin-1beta (IL-1β), Interleukin-10 (IL-10), Interleukin-1alpha (IL-1α) and Tumor Necrosis Factor-alpha (TNF-α)) and chemokines (chemokine (C-X-C motif) ligand 1 (CXCL-1), Monocyte Chemoattractant Protein-1 (MCP-1), chemokine (C-X-C motif) ligand 5 (CXCL-5), Macrophage Inflammatory Protein-1alpha (MIP-1α), Macrophage Inflammatory Protein-1 beta (MIP-1β) and eotaxin) in BALF and plasma were quantified by multiplex beads-based immunoassay technique (LEGENDplex; BioLegend, San Diego, USA) according to manufacturer's instructions. Samples were analyzed with the FACS Canto II (Becton, Dickinson and Company, Franklin Lakes, USA) and data were analyzed with the LEGENDplex data analysis software (BioLegend, San Diego, USA).

**Quantitative polymerase chain reaction (qPCR)**

Lung tissues with Trizol reagent (Invitrogen; Thermo Fisher Scientific, Waltham, USA) were homogenized with Gentle MACS M tubes (Miltenyi Biotec, Bergisch Gladbach, Germany). The total RNA was extracted with RNA purification kit (ZYMO RESEARCH, Irvine, USA) according to manufacturer's instructions. cDNA was synthesized with High-Capacity Reverse Transcription Kit (Applied Biosystems; Thermo Fisher Scientific, Waltham, USA). cDNA was amplified and measured with SYBR Green Master mix (Thermo Fisher Scientific, Waltham, USA) on a CFX96 real-time PCR detection system (Bio-Rad Laboratories, Hercules, USA). Results were normalized to average expression of GAPDH. Target mRNA levels of NV-PBS mice were taken as the control group. Fold change to the control (ΔΔct) was calculated and data were analyzed using the $2^{\Delta\Delta ct}$ method. PCR primers were commercially synthesized (QuantiTect; QIAGEN, Germantown, USA).

**Histology**

Lung tissue for histological examination was obtained from independent experiments as previously described (1). Briefly, tracheal ligation was performed to avoid alveolar collapse. Harvested lungs were fixed in 4% paraformaldehyde solution, paraffin embedded and cut into 2-µm-thick sections followed by dewaxing, dehydration and hematoxylin and eosin (H&E) staining. Histological scoring of lung injury included character, severity, and distribution. All scoring parameters were rated as 0: nonexistent, 1: minimal, 2: mild, 3: moderate, 4: severe. Histopathology examination was performed by a board-certified pathologist, who was blinded to the study groups.

**Electron microscopy**

Lung fixation, subsampling in small tissue blocs of approximately 1 $mm^3$ size and embedding was performed as described previously (6), including a post-fixation with osmium tetroxide, followed by an overnight incubation with uranyl acetate and sample dehydration with ascending acetone concentrations before embedding in epoxy resin. Ultra-thin sections of approximately 70 nm thickness were then cut and images were recorded at a 13000 magnification with a Zeiss Leo 906 TEM (Carl Zeiss, Jena, Germany). Per animal, approximately 70-80 images were randomly recorded of the air-blood barrier and analyzed for integrity of basement membrane.

**Isolated perfused mouse lung ventilation and imaging**

C57BL/6J mice were euthanized (200 mg/kg ketamine, 10 mg/kg xylazine i.p.), intubated via tracheotomy, and connected to the ventilator (HSE mini vent; Hugo Sachs Elektronik - harvard Apparatus GmbH, March-Hugstetten, Germany). Lungs were perfused with Hanks' balanced salts solution (HBSS) (Thermo Fisher Scientific, Waltham, USA) and 4% bovine serum albumin (BSA) (Sigma-Aldrich, St. Louis, USA) *via* the pulmonary artery and ventilated with HVt MV (20 ml/kg) or LVt MV (8ml/kg) for 2h (7). Subsequently, lungs were harvested and 50 µl of the pH sensing fluorescent probe pHrodo™ Red Dextran (0.2 µg/µl; Thermo Fisher Scientific, Waltham, USA) was intratracheally instilled using a micro-sprayer (Penn-Century, Wyndmoor, USA), In every experiment, the patency of airways was checked by changing inflation pressure to exclude injury of the conducting airways and ensure that alveoli expand. The substance delivery by micro-sprayer was previously established in our lab by Gutbier et al. (8), and guaranties that pHrodo™ is uniformly delivered to the lung alveoli. Subpleural alveoli were imaged *in situ* by confocal fluorescence microscopy (Upright Spinning Disk confocal CSU-X / Nikon Ti2; Nikon, Tokyo, Japan) at constant positive airway pressure (CPAP:18 mmHg) monitored by Powerlab data acquisition system (AD INSTRUMENTS, Dunedin, New Zealand).

Images were acquired at 25X magnification using NIS-Elements AR V.5.21.02 software (Nikon; Tokyo; Japan). Nine images from 3 independent experiments per group were analyzed. Integrated fluorescence intensity of randomly selected alveolar sections (non edema filled alveoli) of intermediate size (500-700 μm$^2$) was analyzed with Fiji software (ImageJ, Bethesda, USA).

***In vitro* cell stretch and infection**

Human A549 cells (ATCC #CCL-185) cultured in 2ml medium (RPMI-1640; Gibco; Thermo Fisher Scientific, Waltham, USA) or human primary alveolar epithelial cells (hpAEC; #H-6053, Cell Biologics, Chicago, USA) cultured in respective complete human epithelial cell medium with provided supplements (Cell Biologics, USA) were subjected to cyclic stretch (CS: elongation 18%, 20 cycles/min) for 24h using the FlexCell system (Flexcell PlusTM FX-5000T; Flexcell International Corporation, Burlington, USA). No-cyclic stretch (NCS) control plates were kept in static condition. A pH-electrode (METTLER TOLEDO, Columbus, USA) was used to measure the medium pH. LDH concentration as surrogate for cell death was determined via ELISA. Cells and culture supernatants from CS and NCS conditions were collected for subsequent infection experiments. Accordingly, cells were seeded onto ibidi μ-slides (ibidi GmbH, Gräfelfing, Germany) combined with respective supernatants as indicated in **figure 6A**. After attachment, cells infected with green fluorescence protein (GFP) encoded *P. aeruginosa* PAO1 strain (laboratory stock) at multiplicity of infection (MOI) of 10 for 4h. After infection, cells were thrice washed and stained with following dyes: Alexa Fluor 546 Phalloidin (Thermo Fisher Scientific, Waltham, USA), GFP Polyclonal Antibody Alexa Fluor 488 (Thermo Fisher Scientific, Waltham, USA) and 4′,6-Diamidino-2-phenylindole dihydrochloride (DAPI) (MERCK KGaA, Darmstadt, Germany). Bacterial cellular attachment was assessed by immunofluorescence microscopy (LSM 780; Carl Zeiss Microscopy GmbH, Jena, Germany). Ten (3x3) tile scans were acquired from each condition at 63X magnification. Bacteria to cell ratio was determined using Fiji software (ImageJ, Bethesda, USA).

To determine the pH induced alterations on bacterial fitness, PAO1 growth in acidified, untreated or alkalized RPMI medium was assessed every 30 min by measuring optical density (OD) at 600 nm. HCl and NaOH were used to prepare acidified or alkalized conditioned medium, respectively. In addition, PAO1 was incubated with A549 cells at MOI of 0.1, 1, 5 and 10 in respective pH-conditioned media for 24h and bacterial counts were determined.

To investigate the effect of sodium-hydrogen antiporter 1 (NHE-1) and cystic fibrosis transmembrane conductance regulator (CFTR) on the airway surface pH, A549 cells were incubated with the NHE-1 inhibitor cariporide (50µM; MERCK KGaA, Darmstadt, Germany), CFTR inhibitor - 172 (50µM; MERCK KGaA, Darmstadt, Germany) and dimethyl sulfoxide (DMSO; MERCK KGaA, Darmstadt, Germany) solvent control, respectively. Both cariporide and CFTR inhibitor -172 were dissolved in DMSO and diluted with PBS to a desired concentration. Equal amount of DMSO was used as a solvent control. After 24h of incubation, pH of cell culture medium was determined and cells were then infected with PAO1 at the MOI of 10 for 4h and bacterial counts were determined.

**Data analysis**

Statistical analysis was performed using the GraphPad Prism 9.30 (San Diego, USA) software. Data are presented as mean with SEM or box-and-whisker plots depicting median, quartiles, and range excluding outliers (open circles). CFU data were logarithmized. For comparison of two groups, Wilcoxon matched-pairs signed rank test or Mann-Whitney U test was used and for multiple comparisons of data that was not nonnormally distributed (histological scoring) a two-tailed Mann–Whitney U tests followed by Bonferroni correction was used. For multiple comparisons of normally distributed data as tested by Shapiro-Wilk test, one-way or two-way ANOVA with Sidak's multiple comparisons test was used. P values < 0.05 were considered statistically significant.
